# The structure and evolution of sex chromosomes in an androdioecious species with two mating types

**DOI:** 10.64898/2026.11.09.764214

**Authors:** Adrián Contreras Garrido, Marie Monniaux, Jean Watelet, Ayush Valecha, Anaïs Backland, Céline Poux, Cécile Godé, Clément Mazoyer, Sylvain Legrand, Mathilde Dufay, Philippe Vernet, Xavier Vekemans, Sylvain Billiard, Pierre Saumitou-Laprade, Vincent Castric

## Abstract

Though flowering plants are ancestrally hermaphroditic, some species have evolved sexual specialization and are now composed of individuals expressing just one of their two reproductive functions (either male or female) together with co-sexual individuals. Androdioecy, the co-existence of males with hermaphrodites, is rare across the living world, but unusually common in species of the Oleaceae plant family. The mediterranean shrub Phillyrea angustifolia has a simple mendelian XY sex-determination system for androdioecy, but the underlying molecular determinants and the evolutionary history of the X and Y chromosomes remain unknown. The ability of males to by-pass the self-incompatibility phenotype expressed in hermaphrodites is essential to explain their high frequency in natural populations, but how the loci governing sex and self-incompatibility epistatically interact also remains unknown. To address these questions, we obtained full haplotype-resolved de novo chromosomal assemblies of P. angustifolia and its sister species P. latifolia. The sex-determining region in P. angustifolia spanned a fifth of chromosome 12, and was strikingly larger and more rearranged than that in P. latifolia. The male-specific region contains two putative sex-determining genes, ANDY1 and ANDY2, a tandem encoding two flowering-related transcription factors that arose by translocation followed by duplication of an autosomal gene before the two species diverged. Strikingly, this sex-determining region also contains a recent duplicate of the pollen allergen gene Ole-e1, which shows the strongest differential expression between the two self-incompatibility groups of hermaphrodites, hence establishing a direct molecular link between sex and self-incompatibility. Overall, our results provide novel insight on sex determination and the evolution of sex chromosomes in a plant lineage where the separation between sexes is still incomplete, and offer a possible mechanistic basis for the strong epistatic interaction between the sex and self-incompatibility determinants, with direct implication for how sex-ratios are established in nature.

## Introduction

Flowering plants are ancestrally hermaphroditic (Sauquet et al. 2017), and the vast majority of species (about 85%, (Renner 2014)) have retained the expression of combined male and female sexes by the same (co-sexual) individuals. Yet separate sexes have evolved many times independently, and approximately 6% of known angiosperms are now dioecious (Charlesworth and Charlesworth 1985); (Renner 2014); (Käfer et al. 2017). Several theoretical scenarios have been proposed for the emergence of separate sexes and sex chromosomes from ancestral hermaphroditism. These scenarios involve either the gradual differentiation of increasingly specialised male and female individuals (Lesaffre et al. 2024) or the stepwise invasion of reciprocal sterility mutations directly establishing males and females into the population (Charlesworth and Charlesworth, 1978). In most lineages, however, the genetics of sex determination is unknown and the discovery of sex chromosomes in plants is still at an early stage, with a broad diversity of molecular functions involved (Marais et al. 2025). Beyond the full separation of sexes, plants exhibit a spectacular diversity of sexual polymorphisms, notably including situations where individuals genetically specialized toward either the female or the male reproductive functions coexist with hermaphrodites. The former (gynodioecy) is relatively common and well understood as a result of genetic conflict between nuclear and cytoplasmic genes ((Renner 2014)), but the latter (androdioecy), in contrast, is an exceedingly rare sexual polymorphism across the tree of life ((Renner 2014)), and to the best of our knowledge, the molecular determinants of androdioecy have not been resolved in any plant lineage so far.

In sharp contrast with its extreme rarity in most lineages, androdioecy evolved in the Oleaceae family on several independent occasions (Wallander 2008), providing a promising model system to investigate its genetic basis and evolution. The Oleaceae is an economically important family of plants (*e.g.* the olive tree) and includes the widespread mediterranean shrub *Phillyrea angustifolia*, whose mating system has been extensively studied over the last 30 years (Lepart and Dommée 1992; Francq et al. 2023). A remarkable feature of natural populations of this species and of its sister lineage *P. latifolia,* is that a large fraction of individuals (referred to as “males” below) produces flowers with a small, reddish-brown and cankered stigma (Fig. 1A) that degenerates at late stages of pistil formation. This fully abolishes their female reproductive function, as these individuals bear no fruit. In contrast, hermaphroditic individuals develop perfect flowers with functional male and female organs, and bear fruits (Fig. 1A). In *P. angustifolia*, sex determination follows a XY system, whereby hermaphrodites are homogametic (XX) and males heterogametic (XY), (Billiard et al. 2015)) and the sex-determining region (SDR) is genetically associated with a series of genetic markers to a single linkage group (LG12, (Carré et al. 2021). However, the physical extent of the SDR along the chromosome, whether it exhibits the classical hallmarks of sex chromosome evolution (*e.g.* successive rearrangements, gene loss and repeat accumulation (Saunders and Muyle 2024) and the identity of the genetic elements hindering proper development of the female apparatus and thus determining the male sex in natural populations remain to be discovered.

**Figure 1.**
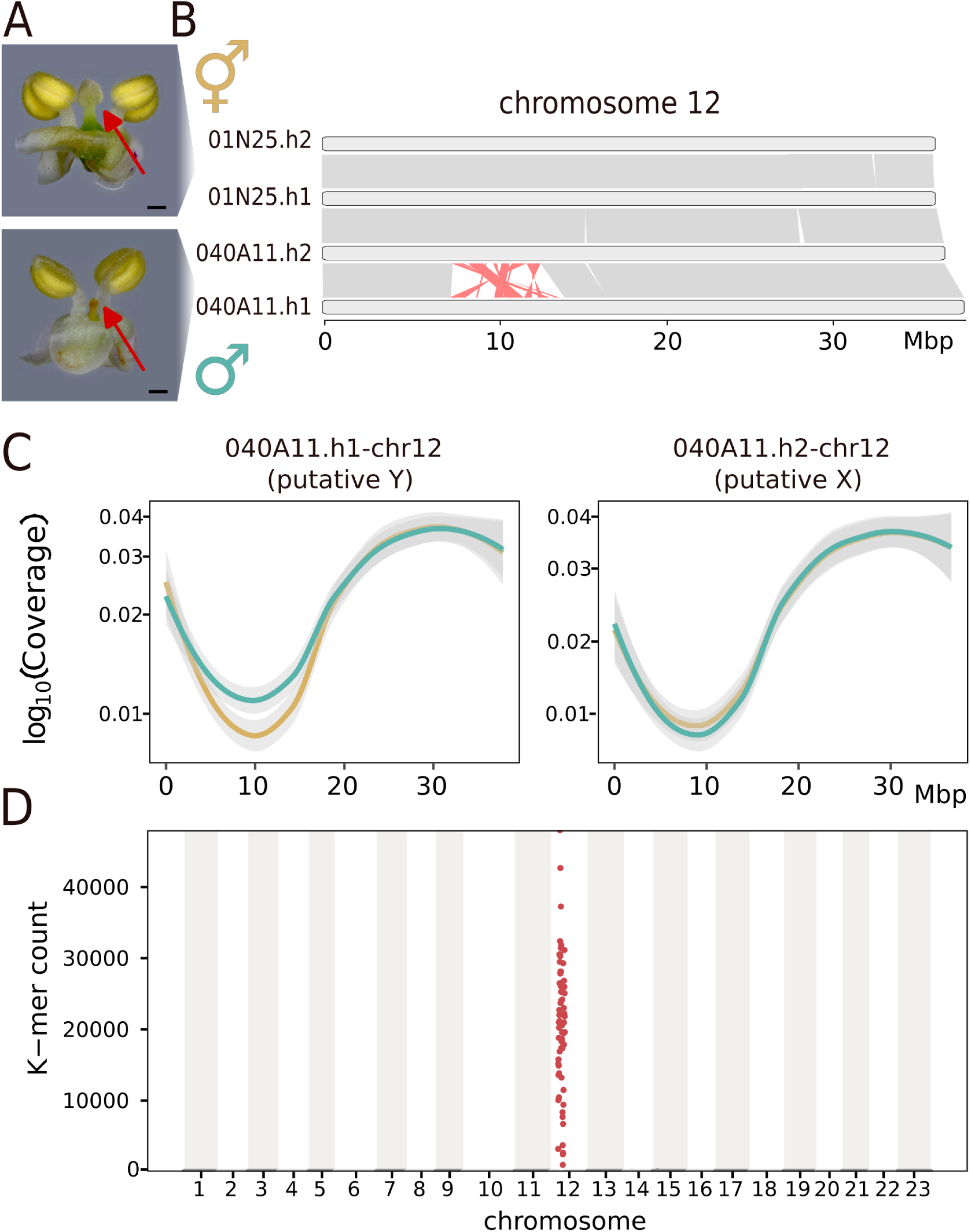
Mapping the *Phillyrea angustifolia* sex determining region. **A)** In *P. angustifolia*, males are distinguished from hermaphrodite individuals by the presence of an aborted stigma in the flowers they produce. The upper panel shows a hermaphrodite flower with a red arrow pointing to its functional stigma; the lower panel shows a male flower with a red arrow pointing to the dysfunctional stigma. Black bar denotes the scale: 0.5 mm. **B)** Structural variation (SV) of chromosome 12 between haplotype 1 of the male genome assembly (040A11.h1) and the three other haplotypes (040A11.h2, 01N25.h1 and 01N25.h2) from the hermaphrodite and the male assembly. **C)** Coverage differences of mapping the published GBS data (Carré et al. 2021) to the two different haplotypes of the male genome assembly (040A11) between males and hermaphrodites of a full-sib controlled cross of *P. angustifolia* shows a cosegregation of the SV with sex phenotype. The green and gold lines correspond to males and hermaphrodites, respectively. Coverage values were fitted using the “gam” method and the grey area corresponds to the 95% confidence interval of the fitted line. **D)** Manhattan plot showing the counts of mapped Y-mers in 100kbp windows along the male assembly (040A11.h1). Y-mers were identified based on whole genome sequencing of 11 males and 10 hermaphrodites from the Fabrègues natural *P. angustifolia* population.

Many lineages within Oleaceae share a particular genetic self-incompatibility system (Saumitou-Laprade et al. 2010) in which hermaphrodites fall into two groups of reciprocal reproductive compatibility, referred as [Ha] and [Hb] hermaphrodite groups (Vernet et al. 2016). While pollen produced by hermaphrodites is compatible with only half of the available pistils in their species (those of their opposite group of compatibility), pollen produced by *P. angustifolia* males are able to fertilize all pistils, regardless of their genotype at the self-incompatibility locus, providing them with an immediate two-fold reproductive advantage over hermaphrodites ((Saumitou-Laprade et al. 2010)). This ability to by-pass the self-incompatibility system is key to the maintenance of males, but the mechanisms of this epistatic suppression remain elusive. In addition, while balanced sex-ratio of the offspring is observed when male pollen is deposited on pistils of [Ha] hermaphrodites (as expected under normal mendelian segregation of X and Y chromosomes), full transmission distortion toward the male phenotype is consistently observed on pistils of the other group of hermaphrodites [Hb] (100% males, 0% hermaphrodites, consistent with selective fertilization by Y-bearing pollen only, (Billiard et al. 2015). Hence, male pollen interacts with pistils of the two groups of hermaphrodites in contrasted manners, representing another level of epistasis between the sex and SI-determining loci. (Castric et al. 2024) recently showed that the two SI groups expressed by hermaphrodites of *Phillyrea* are determined by the presence or absence of a gene predicted to be involved in catabolism of Gibberellin hormones (*GA2ox*), located on chromosome 18, and thus genetically independent from the SDR (Carré et al. 2021). The two groups of hermaphrodites exhibit a number of differentially expressed genes with diverse molecular functions throughout their genome (*n*= 54 in (Castric et al. 2024)), and we hypothesized that exploring the gene content of the SDR may help understand the molecular nature of the epistatic interactions between sex and SI.

Here, we first resolved the genetic architecture and gene content of the SDR of *P. angustifolia,* and showed that it consists of a heavily rearranged ∼6 Mbp region showing the typical hallmarks of a proto sex chromosome, with a decrease in gene content, an increase in transposable elements and putative signs of the progressive cessation of recombination under the form of “evolutionary strata”. Further comparison with *P. latifolia*, a sister species sharing the androdioecy mating system with *P. angustifolia,* indicated that, at its core, the SDR consists of a genomic region that comprises two genes encoding MYB transcription factors of the G2 family. Moreover, examination of the genes within the SDR expressed in flower buds revealed the presence in both species of paralogs of genes of the pollen allergen family *Ole e1*, that rank among the top differentially expressed genes between the two hermaphrodite incompatibility groups. Overall, beyond discovering a novel sex determination mechanism in plants, our study provides insights into an intriguing epistatic interaction between sex and self-incompatibility in an androdioecious species.

## Results

### The heterogametic XY sex determining region of *P. angustifolia* is a heavily rearranged, megabase-sized chromosomal region

To resolve the genetic architecture of the SDR of *P. angustifolia,* we first leveraged the whole chromosome haplotype-resolved assemblies of one male (040A11) and one hermaphrodite (01N25) individuals (Fig. 1A) from our previous work (Castric et al. 2024). Since males are the heterogametic sex (Billiard et al. 2015; Carré et al. 2021), we performed pairwise comparisons among the four haplotypes of chromosome 12, three of which are predicted to carry the ‘X’ allele and one the ‘Y’ allele. These comparisons revealed a highly rearranged region in haplotype 1 of the male genome (040A11.h1) relative to the other three haplotypes (040A11.h2, 01N25.h1, and 01N25.h2) (Fig. 1B), including two rearrangements exceeding 1 Mb and eight exceeding 100 Kb (Fig. S1) from a total of at least 162 rearrangements. This region spanned 5.18 Mb in the X-carrying haplotype (040A11.h2) and 6.71 Mb in the Y-carrying haplotype (040A11.h1). To verify that this rearranged region indeed segregates with sexual phenotypes, we first re-analyzed the Genotyping By Sequencing (GBS) data from the 196 full-sib offspring of the genetic mapping cross (Carré et al. 2021), where sex segregates as a simple mendelian trait. We aligned the GBS sequences to the two haplotypes of the male assembly (040A11.h1 and 040A11.h2) and observed a change in coverage in males *vs.* hermaphrodites consistent with the expected segregation patterns of an XY system (Fig. 1C). Coverage of GBS markers around the rearranged region of the 040A11.h1 haplotype dropped considerably for hermaphrodites, and about half of that drop was observed for males (Fig. 1C), identifying this haplotype to be the Y-carrying chromosome. The opposite pattern was observed when mapping the same GBS sequences to the other haplotype (040A11.h2; Fig 1B), thus corresponding to the X chromosome. No such discrepancy of coverage was observed between the two sexual phenotypes in any other genomic region (Fig S2). The 22 GBS markers that showed full linkage to sex phenotypes in (Carré et al. 2021) were all located either within or in close proximity to the putative SDR in both haplotypes (Fig. S3).

To verify that this structural variant associates with the SDR beyond a single cross, we then examined its presence in short read sequences of ten hermaphrodites and eleven males from a natural population of *P. angustifolia* (Fabrègues, France, see Methods). We extracted K-mers that were common to all male individuals and absent in any hermaphrodite individuals and mapped them to the assembly containing the putative Y-SDR. 99% of them mapped to the structural variant in chromosome 12 observed in the assembly (Fig 1D; 2A). We also measured the levels of genetic differentiation (using Weir & Cockerham’s *F*_ST (Weir and Cockerham 1984)_) considering males and hermaphrodites as separated populations and observed high differentiation at the putative SDR (Fig. 2A). Taken together, our analyses confirm that the haplotype-specific structural variant found in the male assembly 040A11 corresponds indeed to the SDR of *P. angustifolia*, and that the SDR spans about a fifth of the length of chromosome 12, the rest constituting two large pseudo-autosomal regions (PARs).

**Figure 2.**
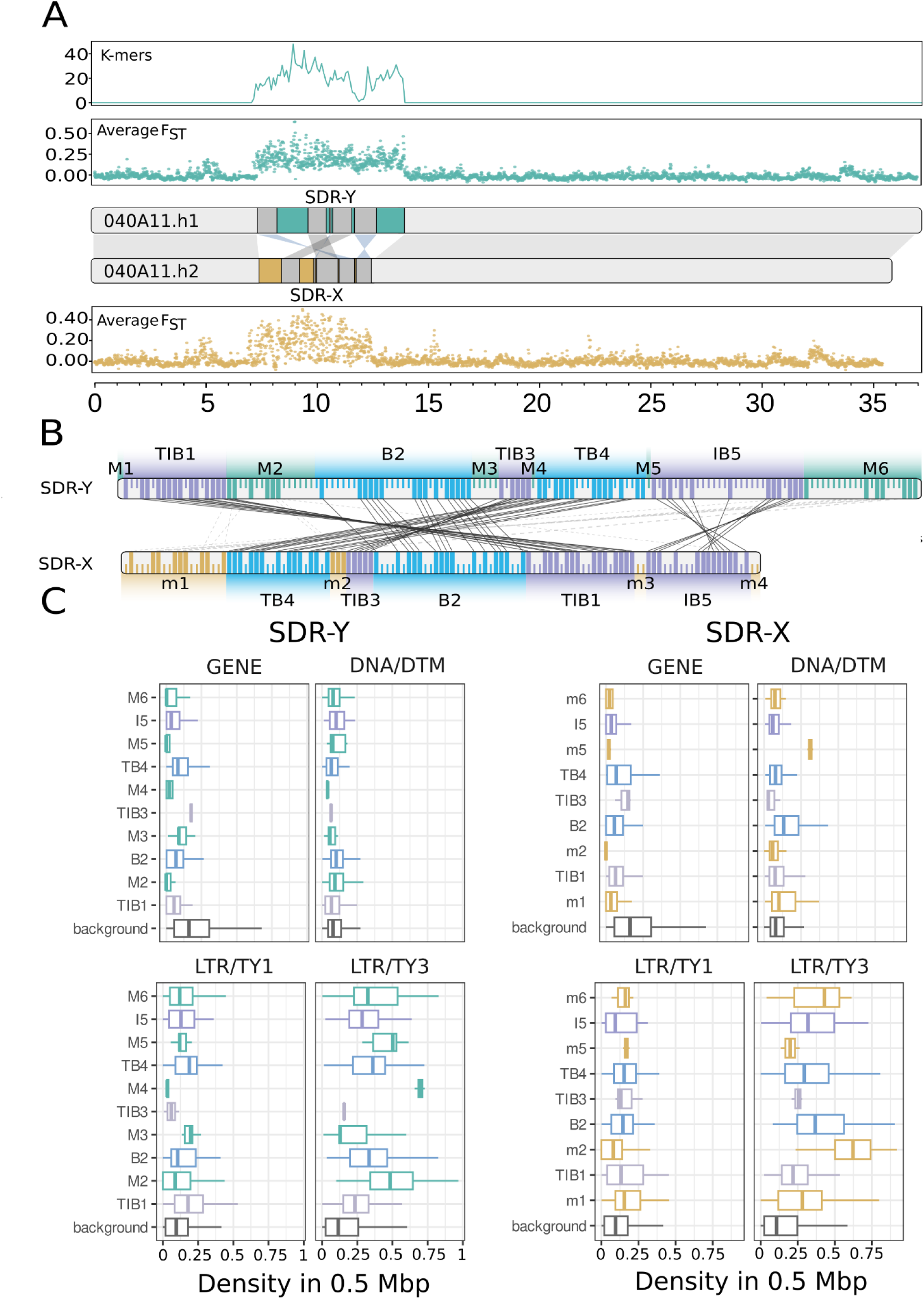
*Phillyrea angustifolia* SDR composition. A) Chromosome-scale representation of the SDR and population-level evidence of its localization on chromosome 12. The upper panel shows overlap between k-mers found exclusively in male samples, mapped against the Y-bearing haplotype (040A11.h1) and the SDR structure in the third panel. The second and fourth panels display genetic differentiation between males and hermaphrodites, measured as Weir and Cockerham’s *F*_ST_ (Weir and Cockerham 1984), along haplotypes 040A11.h1 and 040A11.h2, respectively. The third panel illustrates the syntenic relationships between the two haplotypes of 040A11, as inferred by GENESPACE. B) Relationships between genes in the SDR regions of the X and Y haplotypes, 040A11.h1 and 040A11.h2, respectively. Genes are ordered sequentially from 5’ (left) to 3’ (right) along each haplotype, but actual physical distance is not represented. Homologous genes are represented by long ticks of each haplotype and connected by lines between haplotypes. Black lines indicate homologous genes that remain in synteny between haplotypes, whereas grey dotted lines indicate homologs that are rearranged between the haplotypes. Genes unique to either the X or Y SDR, private genes, are shown as short ticks on the corresponding haplotype. Tick color as well as background color denotes the syntenic block to which each gene belongs to. C) Density analysis for different genomic features at each synteny block of the SDR and at the rest of the genome, referred to as background, for both 040A11.h1 and 040A11.h2 haplotypes. Density is measured in counts per window of 0.5 Mbp.

### Recombination arrest and the formation of evolutionary strata in *P. angustifolia* SDR

To clarify the structural conformation of the SDR and evaluate the extent of rearrangements between the X and Y alleles, we examined the genic synteny between both haplotypes at the SDR and the rest of the chromosome. We used GENESPACE (Lovell et al. 2022) to compare the four haplotypes derived from the male and hermaphrodite assemblies. There was perfect collinearity among the three X-carrying haplotypes at the SDR, and all of them showed the same distinctive rearrangement pattern when compared with the Y SDR allele (Fig. S4). These rearrangements comprise six non-syntenic blocks that were Y-specific (M1 to M6 on Fig. 2A and 2B) and six that were X-specific (m1 to m6). In addition, five syntenic blocks within the SDR were shared between the X and Y alleles. Among them, only one block (B2) appears to have retained the same position and orientation in both alleles, the others being either inverted (IB5), translocated (TB4) or both (TIB1, TIB3; Fig. 2A and 2B) with respect to one another.

We next examined the gene content of the SDR in *P. angustifolia* between the X and Y SDR alleles. The Y-SDR harbours 146 putative genes, and the X-SDR only 117. Among these, we could identify 72 homologous pairs of genes between the two alleles, including 59 pairs that retained their relative genomic positions (syntenic) and 13 pairs that were rearranged between the X and the Y alleles, indicating that besides the overall high-order syntenic structure highlighted above, small-scale structural rearrangements also occurred within the two alleles, often involving a single gene (Fig. 2B). The remaining genes, 74 on the Y-SDR and 45 on the X-SDR, were classified as private. When correcting for differences in SDR size, gene density was nearly identical between the Y- and X-SDR alleles (0.0806 and 0.0815 genes per Mb), but sharply lower than the background genome-wide average (0.2034 genes per Mb, Fig. 2C). This pattern was mirrored in the overall TE densities, which were higher in the SDR as compared to the genome background in both haplotypes, but were similar between the X- and Y-SDR alleles. In particular, both SDR haplotypes showed an enrichment of long terminal repeat (LTR) retrotransposons relative to the genomic background (Fig. 2C). LTR/Ty3 transposons were twice as abundant in the X- and Y-SDR sequences (∼0.33 elements per Mb at both alleles) as at the genomic background (∼0.16 elements per Mb). A moderate enrichment was also observed for LTR/Ty1 transposons at the SDR (∼0.15 elements per Mb) as compared to the genomic background (∼0.11 elements per Mb). This pattern was not shared across all transposable element classes, however, and DNA Mutator transposons, the most abundant TE family in *P. angustifolia*, showed no appreciable enrichment within either SDR haplotype relative to the genomic background (Fig. 2C).

Interestingly, TEs located within the SDRs tended to be on average younger than those found elsewhere in the genome, as inferred from both LTR sequence similarity and divergence from family consensus sequences. This trend was particularly pronounced for LTR/Ty3 and DNA/DTM transposons, for which the differences were statistically significant (Fig. S5). Overall, contrary to the expectation from classical theory (Charlesworth and Charlesworth 2000), in *P. angustifolia* we do not find greater degeneration (in the form of gene loss and TE accumulation) of the SDR of the Y chromosome as compared to that of the X.

This lack of observable degeneration in the Y allele suggested that the observed SDR configuration might be recent and prompted us to investigate the divergence time between the two SDR alleles. If recombination between X and Y has indeed stopped, then we can use the sequence divergence between the identified homologous gene pairs at the SDR to estimate its age. To do this, we calculated synonymous divergence (dS) for the single-copy syntenic genes within the SDR as well as genome-wide single copy genes across both haplotypes of 040A11 (see Methods). The median synonymous divergence for the SDR alleles was more than three times higher than the genome-wide median between haplotypes (dS=0.0283 vs. 0.0085, *p*<0.0001; Fig. 3A and Fig. S6), supporting that recombination between the two SDR alleles indeed ceased. The mean dS value varied among the different synteny blocks (Fig. 3B), suggesting that recombination stopped at different times in the past, and they may represent successive evolutionary strata (Bergero et al. 2007)(Akagi et al. 2025). Specifically, divergence peaked in the central region of the SDR (block B2) and declined toward the chromosome arms in a stepwise pattern, with the exception of TIB3 (which contained only three shared syntenic genes, leading to a very uncertain estimation). This stepwise structure was further supported by pairwise DNA alignments between the blocks (Fig. S7). In line with this scenario, the accumulation of LTR/Ty3 elements at the syntenic blocks also peaked at B2 (Fig. 2C).

**Figure 3.**
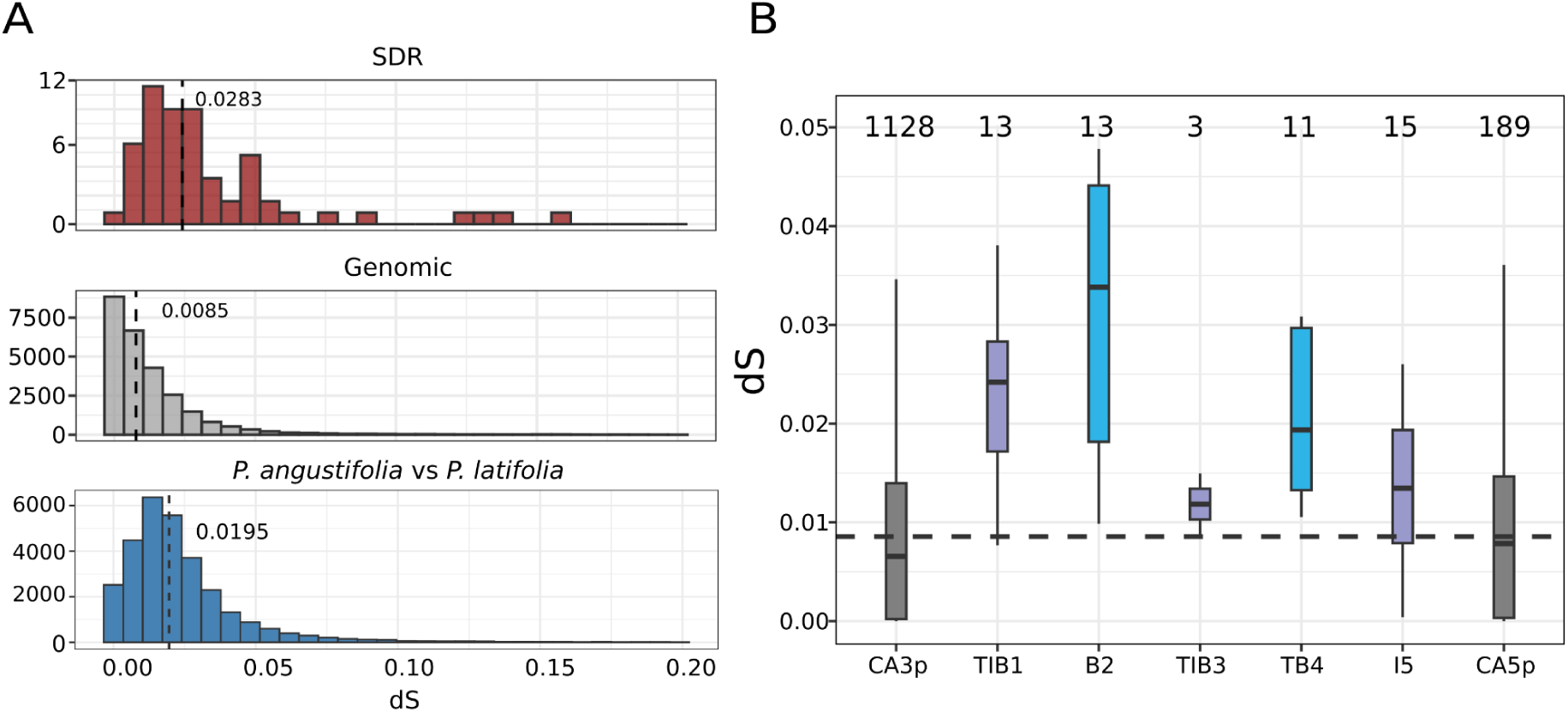
Evolutionary strata of the SDR of *P. angustifolia*. A) Distribution of the overall dS values between haplotypes of 040A11 for genes in the SDR (upper panel); between haplotypes of 040A11 for genes across the rest of the genome (middle panel); between *P. angustifolia* and *P. latifoli*a homologs (lower panel). In all three panels, dotted lines display median values. B) Synonymous divergence (dS) between X and Y alleles at the *P. angustifolia* SDR partitioned by synteny blocks (mute lavender indicates syntenic inverted blocks, blue represent syntenic block that retain the same orientation and grey represent the pseudo-autosomal regions). The dotted line indicates the genome-wide median dS (0.0085). The number of homologous gene pairs from which dS was estimated is shown above each boxplot. CA3p and CA5p refer to the pseudo-autosomal regions chromosome 12 arms on the 3ʹ and 5ʹ sides of the SDR, respectively.

### Most rearrangements in the Y-SDR are not observed in the sister species *P. latifolia*

We then extended our analysis to the sister species *P. latifolia*. Both species share the androdioecious reproductive system, and the male phenotype in *P. latifolia* similarly entails developmental arrest of the stigma (Fig. S9). Furthermore, male pollen of *P. latifolia* also has the property of being able to fertilize both groups of hermaphrodites, and by analysing 118 offspring from two replicate controlled crosses, we found that male pollen of *P. latifolia* also shows massive transmission distortion on [Hb] pistils of *P. angustifolia (*Table S2). Hence, besides the macroscopic similarities of male flowers, in both species the sex and SI systems interact in a similar way. As such, we hypothesized that the molecular determinants of sex and sex-SI interaction should also be shared between the two species. Following this logic, we sequenced and assembled the genome of a male individual of *P. latifolia,* assuming that it should contain both the X- and Y-SDR alleles. The k-mer based estimation of the genome size on the unassembled HiFi reads was 785 Mbp, similar to that of the *P. angustifolia* individuals we previously assembled (Castric et al. 2024). *P. latifolia* assembly (Table S3) also showed a very good contiguity (N50=31.9 Mbp), high completeness (99.3 % of BUSCO genes present) and genome-wide heterozygosity of 1.75%. The mean synonymous divergence of homologous gene pairs between the two species was 0.0195 (n=30,962 homologs), which was slightly lower than the synonymous divergence observed between the two SDR alleles in *P. angustisfolia* (dS = 0.0283; Fig. 3A). This suggests that the X and Y SDR alleles stopped exchanging genetic material through recombination before the split between the two species.

Albeit we observed almost perfect synteny between both species, with a 1:1 relationship between chromosomes of both haplotypes of *P. latifolia* and those from *P. angustifolia* 040A11 (Fig. S9), chromosome 12 stood out as an exception to this overall similarity. While synteny was very good along the chromosome arms, the region containing the putative SDR was an exception (Fig. 4A). One of the two *P. latifolia* haplotypes showed strong synteny across the SDR region with the *P. angustifolia* X-SDR allele, indicating that it likely represents the homologous *P. latifolia* X-SDR allele. Remarkably, the other haplotype lacked synteny with the *P. angustifolia*’s Y-SDR, indicating distinct structural configurations of the Y chromosomes of the two sister species.

**Figure 4.**
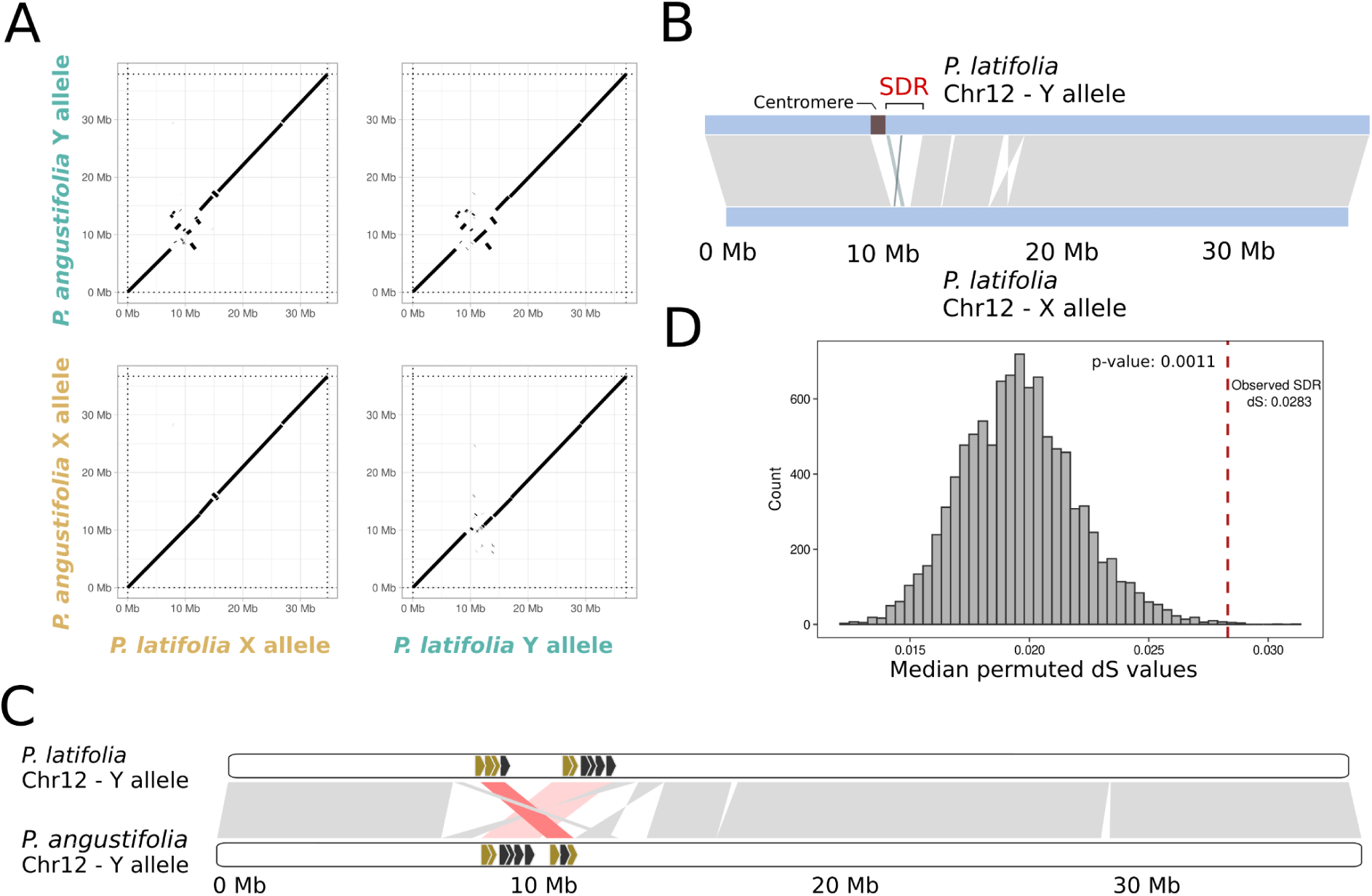
The core Phillyrea SDR. A) Dotplots showing syntenic relationships in chromosome 12 between the haplotypes of sequenced male individuals of *P. latifolia* and its homologous haplotypes in *P. angustifolia. B)* Syntenic alignment between chromosome 12 haplotypes containing Y- and X-SDR loci in *P. latifolia*. C) Syntenic schematic of the comparison of the two Phillyrea Y-SDR alleles, with the approximate positions of the nine conserved genes between the two alleles and, in golden, the final four candidate genes considered for the male syndrome. D) The mean synonymous divergence (dS) between homologous gene copies on the X- and the Y-SDR of *P. angustifolia* (dS=0.0283; dotted vertical line) is significantly higher than that between orthologous genes in *P. angustifolia* and *P. latifolia*, suggesting that the X- and Y-chromosomes stopped recombining before the two species became separated. The null distribution was obtained by 10,000 resampling of 55 genes (the number of homologous genes in the SDR randomly chosen across the dS values of *P. angustifolia* and *P. latifolia* homologs).

Indeed, the *P. latifolia* SDR configuration was simpler than in *P. angustifolia*, with three consecutive non-homologous regions between the X and Y SDR alleles separated by two small inversions (200Kbp and 111 Kbp; Fig. 4B). We excluded the first of the three non-homologous regions due to their overlap with the centromere, considering the rest as the SDR locus of *P. latifolia*. The resulting region was 2.02 Mbp in size for the Y-SDR allele and 1.06 Mbp for the X-SDR, both considerably shorter than their corresponding homologs in *P. angustifolia* (6.71 Mbp for Y-SDR and 5.18 Mbp for the X-SDR). Finally, we compared the gene content of the SDR alleles of the two Phillyrea species. A single gene was shared among the four SDR alleles. Nine were uniquely conserved but inverted between the two Y-SDR alleles (Fig. 4C; Table S4). These rearrangements between the Y-SDR alleles contrast with the perfect collinearity of the X-SDR alleles of the two species (Fig. 4A). We then compared the median dS between the X and Y SDR alleles previously estimated in *P. angustifolia* with the overall genome-wide divergence between orthologous genes of *P. angustifolia* and *P. latifolia*. To account for the disparity in sample sizes, we performed a bootstrap test (see Methods). The divergence between X and Y alleles of *P. angustifolia* was significantly higher than the interspecific divergence (Fig. 4D), indicating that the formation of the X and Y alleles predates the split between the two species. This places the origin of androdioecy before the speciation event of *P. angustifolia* and *P. latifolia*, estimated to have taken place around 8.5 MYA (Dupin et al. 2022).

### The SDR is strongly enriched for genes with differential expression between sex morphs

To narrow down which genes are causal for the male sex phenotype in *P. angustifolia*, we complemented the expression data from 14 hermaphrodites examined in (Castric et al. 2024) with 12 males from the same sequencing experiment but presented here for the first time. These 26 samples were collected from mixed stage flower buds before flowering. To minimize reference bias for genes within the SDR alleles, we mapped the RNA-seq reads on a concatenated version of the male *P. angustifolia* reference genome (040A11), in which both the X- and Y-containing haplotypes are present. We called differentially expressed genes (DEGs) for X- and Y-SDR containing haplotypes separately (see Methods), which allowed us to analyse both sets of differentially expressed genes, in the X and the Y, independently.

Overall, using the Y-containing haplotype (040A11.h1) as a reference, 343 genes were differentially expressed between males and hermaphrodites, 49 of which (14%) were located within the Y-SDR. Similarly, using the X-containing haplotype as a reference (040A11.h2), 288 genes were DEGs between males and hermaphrodites, 31 of which (10%) were located within the X-SDR. Hence, while the *P. angustifolia* SDR represents a relatively small proportion of the total genome (0.69% for the X and 0.88% for the Y) and, as we have seen before, also relatively depleted of genes, it is strongly enriched for DEGs between the two sexual morphs in flower buds (Fig. 5A; Fig. 5B; Fig S11, Table S5). Among the nine genes conserved in both *P. angustifolia* and *P. latifolia* at the Y-SDR (Table S4), five show no expression, whereas the remaining four are strongly expressed in males. Notably, three of these four had the highest log2-fold changes across the entire genome, with the fourth also falling within the top ten DEGs between morphs (Fig. 5C). Therefore, we considered these four genes: Pan-gene-95.2, Pan-gene-95.26, Pan-gene-136.0 and Pan-gene-136.2, that combine a conserved status at the Y-SDR locus in the two androdioecious species with a high and specific expression in male flower buds, as the best candidates for the male phenotype in *P. angustifolia*.

**Figure 5.**
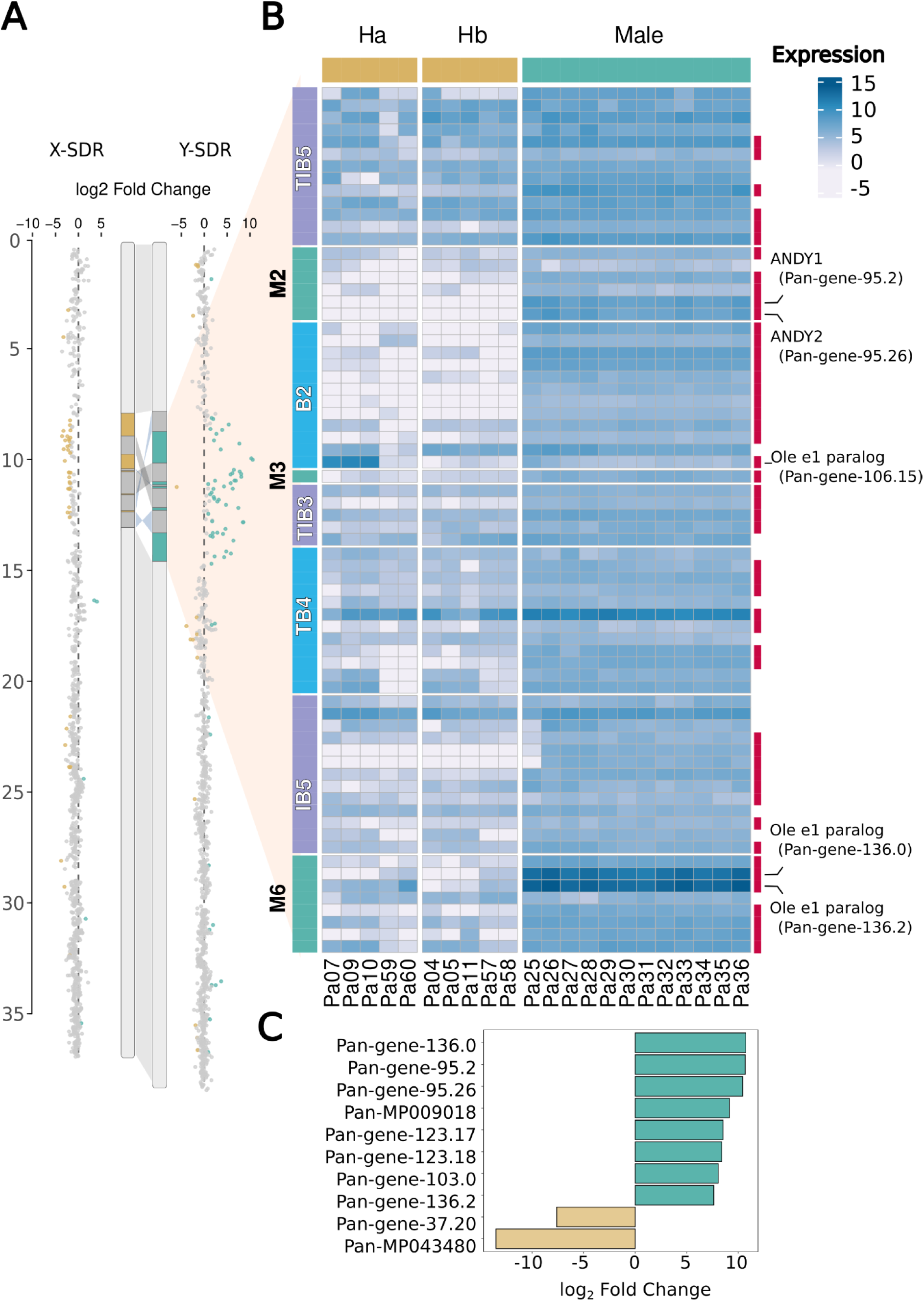
Differential gene expression analysis between males and hermaphrodites in mixed-stages flower buds. **A)** Differential gene expression of genes on chromosome 12. Each dot corresponds to a gene. Gold dots indicate genes with significantly higher transcript levels in males as compared to hermaphrodites (|log2-fold change| > 1 and an adjusted p-value cutoff of 0.01). Golden dots indicate genes with significantly higher transcript levels in hermaphrodites as compared to males (|log2-fold change| > 1 and an adjusted p-value cutoff of 0.01). Grey dots indicate genes whose transcript levels were not detectably different between males and hermaphrodites. **B)** Heatmap of gene expression within the Y-SDR after applying a variance-stabilizing transformation to the count data. Differentially expressed genes are marked by red bars. **C)** Top 10 most differentially expressed genes between hermaphrodites and males in log₂ fold change at 040A11.h1; a vertical red bar denotes which genes are at the SDR.

### Characterization and evolutionary origin of the four candidate genes for sex determination and its epistatic interaction with SI

These four candidate genes consisted of two pairs of gene duplicates. The first pair of genes, Pan-gene-95.2 and Pan-gene-95.26, which we named *ANDRODIOECY Y-LINKED 1 (ANDY1)* and *ANDRODIOECY Y-LINKED* 2 (*ANDY2*) are placed in close physical proximity in the male-specific block M2 (Fig. 2B). They encode transcription factors containing a predicted MYB-like DNA-binding domain (myb_*SHAQKYF;* Pfam PF00249, accession TIGR01557 CD database (Wang et al. 2023)) and are classified as members of the G2 transcription factor family according to the PlantTFDB 4.0 (Jin et al. 2017).

The closest *Arabidopsis thaliana* homolog of *ANDY1/2* is *ULTRAPETALA INTERACTING FACTOR1* (*UIF1*, AT4G37180.1). In *A. thaliana, UIF1* is the only member of this gene family and it was shown to control the activity of floral meristems, playing an essential role in the development of floral organs (Moreau et al. 2016). In spite of the relatively weak similarity to *A. thaliana* UIF1 (∼39% protein identity), both *P. angustifolia* copies retain the key conserved amino acid residues found in UIF1 homologs ((Moreau et al. 2016); Fig. S12). This conservation suggests that the two copies located within the Y-SDR may play an important role in the arrest of pistil development in Phillyrea male flowers. To further sustain the association between the presence of *ANDY1/2* and flower morphs, we designed a genetic marker based on KASPar probes to track presence/absence of these linked genes across a panel of natural accessions of *P. angustifolia* (n=45), and *P. latifolia* (n=19) and found in both species a perfect association between their presence in an individual and the male sex phenotype (Fig. S13, Table S1, Table S7),

To gain insight into the evolutionary origins of the *ANDY* gene pair, we conducted a phylogenetic reconstruction of gene families across Oleaceae. We used publicly available proteomes of several species spanning major clades of Oleaceae: *Abelliophyllum distichum, Forsythia ovata and F. suspensa, Syringa oblata, Fraxinus pennsylvanica, F. excelsior, Olea europaea* and a species from the sister genus of Phillyrea, *Osmanthus fragrans* (Dupin et al. 2024), which is a known androdioecious species. We completed this dataset incorporating the proteomes predicted from the two haplotypes of both *P. angustifolia* and *P. latifolia*. Identification of gene families and the phylogenetic inference into groups of orthologs and paralogs was done using Orthofinder (Emms and Kelly 2019). The phylogenetic relationships between these species is displayed in (Fig. 6A).

**Figure 6.**
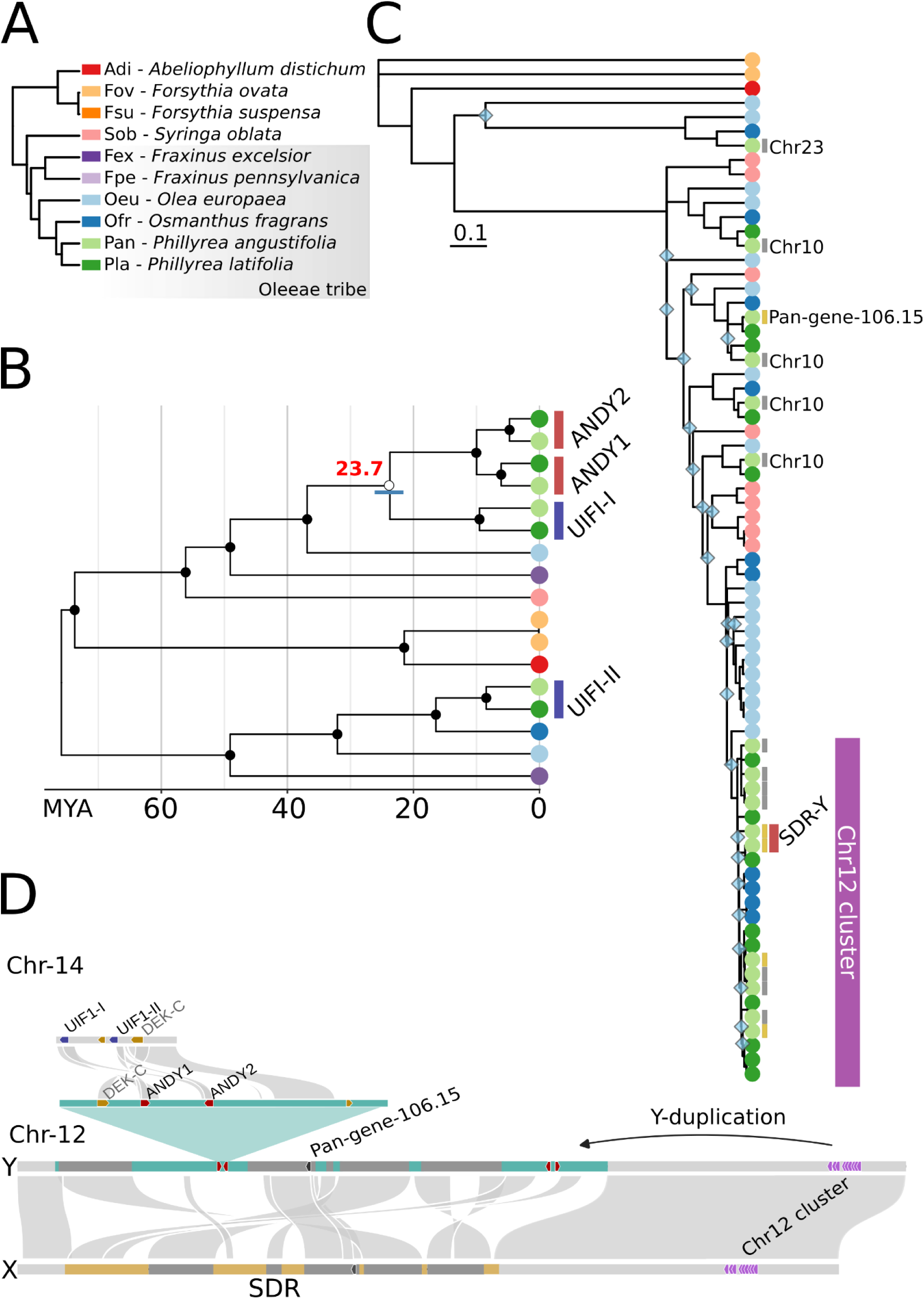
A) Cladogram of the Oleaceae species included in the phylogenetic analyses. Species relationships follow (Dupin et al. 2024). (B) Chronogram of the UIF paralog gene family. Tips are colored according to species, following the color scheme shown in A. The estimated timing of the duplication event that gave rise to the ANDY1/2 clade from UIF1-1 is indicated by a red number. The blue bar represents the uncertainty interval for this estimate, ranging from 21.5 to 26.1 million years ago (MYA), X axis represents divergence time in millions of years. Nodes used for calibration are represented on Fig. S14. Black dots at each node indicate >99% bootstrap support. C) Maximum-likelihood phylogenetic trees of the Ole e1 gene family. The inferred tree was transformed into an ultrametric tree such that all terminal taxa are aligned at the present; the scale bar represents relative evolutionary time after ultrametric transformation. Blue diamonds at nodes indicate >70% bootstrap support. Tips are colored according to species, following the color scheme shown in A. Vertical bars indicate expression differences in *P. angustifolia* during flower development between hermaphrodites and males as previously reported in this work. Grey bars denote genes that are not differentially expressed, whereas gold bars denote differentially expressed genes. The red bar denotes Ole e1 orthologs placed in the Y-SDR allele. D) Proposed model illustrating the origin of Y-linked genes on the Phillyrea Y chromosome through transposed gene duplication. In red, Y-specific genes that arose from a Phillyrea private duplication, displayed here and in B) and C).

Like in *A. thaliana*, the outgroup *A. distichum* contains a single copy of the putative *UIF1 ortholog*, while *F. ovata* has undergone a very recent duplication with two nearly identical paralogs (Fig. 6B). In contrast, species of the Oleeae tribe together with *S. oblata* exhibit at least two deeply divergent copies, *UIF1-I* and *UIF1-II*, each forming a clear phylogenetic clade across species, suggesting an ancestral duplication dating back to the origin of the tribe, about 76 Myrs ago (Fig. 6B; (Dupin et al. 2024)). In *P. angustifolia* these two paralogs are adjacent to one another on chromosome 14, suggesting that they resulted from an ancient tandem duplication. The additional male-specific paralogs *ANDY1/2* are specific to the genus Phillyrea, and their phylogenetic placement indicates that they both derive from *UIF1-I*, rather than from a segmental duplication of both copies together, at a time estimated between 21.5 and 26.1 million years ago (Fig. 6B). The divergence between Phillyrea and Osmanthus dates back to 23 Myrs ago (Dupin et al. 2022), but interestingly, we could find neither *ANDY1/2* nor *UIF1-I* in the raw genomic reads of a sequenced male individual of the androdioecious species *Osmanthus fragrans (Yang et al. 2018)* (Fig. S15, see Methods), possibly indicating a different mode of control of androdioecy in that lineage. Overall, this points towards a scenario where a translocation followed by a duplication of *UIF1-I* gave rise to the *ANDY-1* and *ANDY-2* pair. This duplication event led to the developmental arrest of the stigma of a subset of individuals in the ancestrally hermaphroditic species, abolishing their female function and causing them to reproduce exclusively as males. Further work will be necessary to determine through which pathway the expression of these genes mediates developmental arrest of the stigma, and whether both *ANDY-1* and *ANDY-2* are necessary or play redundant roles.

The second pair of conserved genes located in the male-specific block M6 consists of two genes encoding pollen proteins of the Ole e1 family “*Pollen protein Ole e 1 like; Pfam PF01190*”. These proteins are known as major allergen proteins across the Oleaceae family (Martín-Orozco et al. 1994), with extremely high abundance in pollen (Rodríguez et al. 2002) and they likely play a role in recognition processes between pollen and stigma, as well as between pollen tubes and stylar cells (de Dios Alché et al. 2004; Rodriguez-Rajo et al. 2010). In *P. angustifolia,* a set of 15 *Ole e1* protein-coding genes rank among the top differentially expressed genes between the [Ha] and [Hb] SI specificities in hermaphrodites (Castric et al. 2024). It is remarkable to find two paralogs of this exact gene family conserved in the Y-SDR allele, especially given the ability of male pollen to by-pass the SI reaction.

As mentioned before, Ole e1 transcripts are differentially expressed between the two self-incompatibility types ([Ha] and [Hb]), with higher expression in [Ha] individuals (who lack the GA2ox gene) (Castric et al. 2024). In this work, we observe that the expression of most *Ole e1* paralogs in males resembled that in [Ha] rather than [Hb] hermaphrodites, *i.e.* it was similarly high in [Ha] and males (Fig. S16). Because [Hb] individuals express GA2ox in their stigmatic tissue, which males lack regardless of their SI type, we postulate that GA2ox expression in stigmatic tissue suppresses *Ole e1* expression. The high expression of *Ole e1* genes in [Ha] and males would then be caused either by lack of the *GA2ox* gene (absent from the genome of [Ha] hermaphrodites), or by lack of the tissue where this gene would normally be expressed (in males regardless of their S-locus genotype).

Additionally, the two *Ole e1* conserved paralogs found exclusively in Phillyrea males are also highly expressed in flower buds, resulting in extreme levels of total *Ole e1* transcripts in males beyond that of either [Ha] or [Hb]. Whether and how this extreme composition of male pollen in a key component of the SI phenotype (Ole e1) positively or negatively affects the response they eventually trigger on [Ha] and [Hb] pistils is unknown at the moment, but the mere presence and expression of two *Ole e1* paralogs precisely at the sex locus strongly suggests that they likely play a prominent role in the way sex determination interferes with expression of the SI system.

Our phylogenetic analysis of the *Ole e1* paralogs revealed a single member of this gene family in the outgroups Forsythia and Abeliophyllum, but a series of repeated parallel expansions in all species of the Oleeae tribe. As a result, *P. angustifolia* contains six *Ole e1* paralogs outside of chromosome 12 (five of which are present in *P. latifolia* as well), together with a cluster of nine paralogs arranged as a tandem array in the pseudo-autosomal region of chromosome 12 (eleven paralogs in this cluster in *P. latifolia*). The Y-SDR region of *P. angustifolia* has two *Ole e1* paralogs in the Y-specific M6 region (Pan-gene-136.0 and Pan-gene-136.2; both highly expressed in males, Fig. 5B) and one in the B2 block shared between the X- and the Y-SDRs (Pan-gene-106.15; highly expressed in [Ha] hermaphrodites but not in males, Fig. 5B, S16). Similar to the ANDY1/2 origin, these duplications are unique to the Phillyrea genus. The poor resolution and low bootstrap support of many nodes of the phylogenetic tree of *Ole e1* sequences prevents reliable estimation of the time when these duplications occurred, but placement of the two Y-specific copies clearly establishes that they originated from the cluster in the pseudo-autosomal region of chromosome 12 (Fig. 6C), whereas Pan-gene-106.15 originated from a duplication of one of the paralogs in chromosome 10 (Fig. 6C).

## Discussion

### A newly evolved pair of genes involved in floral development acts as putative female suppression factors

While sex-determining genes have been identified in some plant species, all cases studied corresponded either to species with full separation of the sexes, hence reciprocal suppression of the alternative sex, or to species where the male function only is suppressed in some individuals (gynodioecy) (Marais et al. 2025). Here, we have identified the first putative female-suppression factor that segregates in a plant species in absence of male sterility factors. The suppression is associated with male-specific transcription factors, ANDY1/2, putatively leading to late developmental arrest of the female apparatus. Interestingly, the male-specific gene *aspTDF1* on the Y chromosome of the dioecious *Asparagus officinalis* shares the same protein domain as ANDY1/2 (Myb_DNA-binding domain: PF00249), suggesting that in spite of the large diversity of molecular functions involved in sex determination identified so far among plant lineages (Marais et al. 2025), there might still be some level of convergence at the molecular level. Indeed, *ANDY1/2* are distant paralogs of Arabidopsis UIF1, a transcription factor shown to repress the expression of the stem cell identity gene *WUSCHEL*, itself a known activator of the stamen and carpel identity gene *AGAMOUS* (Moreau et al. 2016). WUSCHEL and its ligand CLAVATA were already associated with gynoecium suppression in the dioecious plant *Silene latifolia* (Kazama et al. 2022). Therefore, late and indirect repression of *AGAMOUS* by ANDY1/2 is a possible and testable mechanism for stigma developmental arrest in *Phillyrea* male flowers.

### Gene amplification as a source of novelty during plant sex chromosome evolution

Strikingly, while at the morphological level the male phenotype entails loss of proper female development, at the molecular level the mutation we identified does not correspond to a classical loss-of-function mutation. Instead, our phylogenetic analysis indicates that female sterility is associated with neofunctionalization of ANDY1/2, a novel pair of paralogs of the IUF1 gene. In line with this observation, (Marais et al. 2025) identified gene duplication as the most common driver for the emergence of sex-determining genes. While whole-genome duplications have occurred repeatedly throughout angiosperm evolution (Renny-Byfield and Wendel 2014) and have been recurrent during the diversification of the Oleaceae family (Julca et al. 2018), here we find instead that ANDY1/2 became established following a simple duplication from autosomal copies before diversification of the Phillyrea genus (Fig. 6D). As mentioned above, androdioecy is unusually common in species of the Oleaceae family, raising the question of whether the same mechanisms are suppressing the female function in other species. *O. fragrans* is closely related with Phillyrea and also exhibits androdioecy, but we could not find ANDY1/2 in genomic sequences from a male individual, suggesting the possibility that a different mechanism may determine the male sex in that genus. Formally testing this hypothesis will require more detailed genomic investigations. More generally, it will be interesting to expand the scope and assess whether Oleaceae species that exhibit complete separation of the male and female sexes (Wallander and Albert 2000) eventually suppress the female function using a similar pathway, as would be expected if androdioecy was an intermediate stage toward full dioecy, a possibility that has long been considered improbable, if not entirely impossible (Pannell 2018).

### A possible mechanism for the epistatic interactions between sex and SI

A possible *a priori* scenario for the epistatic interactions between sex and SI would have been be that male pollen exhibits [Hb]-like expression by default (explaining their full compatibility on [Ha] pistils), and that their Y-bearing pollen “camouflage” into [Ha] when deposited onto [Hb] pistils through the conditional expression of some Y-linked factor in male pollen. This scenario would have explained the ability of male pollen to fertilize both [Ha] and [Hb] pistils, and their distorted transmission onto [Hb] pistils. The fact that the SDR of the Y chromosome contains paralogs of *Ole e1,* the top differentially expressed gene family between [Ha] and [Hb] makes them particularly promising candidates for such epistatic interaction between sex and SI. However, at the moment our observations do not support this simple model of interaction. Instead, the transcriptional profile of males for genes exhibiting strong differential expression between the two groups of hermaphrodites (*Ole e1*) resembles more that of [Ha] than [Hb] individuals.

While the exact mechanisms by which the novel paralogs ANDY1/2 cause loss of the female function, and those by which the novel *Ole e1* paralogs in the SDR modify expression of the SI phenotype of male pollen remain to be established, establishing more precisely the sequence of events remains challenging. In principle, males cannot be maintained in a population unless they are able to compensate for the loss of their female function. Hence, one would expect that the *Ole e1* paralogs arose first, creating a transient class of hermaphrodites with universal compatibility. Such a situation was indeed reported in the Oleaceae *Ligustrum vulgare*, where hermaphrodites with universally compatible pollen segregate at a low frequency in natural populations (De Cauwer et al. 2021). Under this scenario, subsequent loss of the female function by acquisition of the ANDY1/2 paralogs would then allow these hermaphrodites to avoid producing selfed progeny suffering from the deleterious effects of inbreeding depression. Testing this scenario will be an exciting next step for future research that will require a better understanding of the complex diversification history of the *Ole e1* gene family.

### Contrasted patterns of chromosomal degeneration between sister species of Phillyrea

Rearrangements are a classical hallmark of sex chromosome evolution, and are classically interpreted as the consequence of the linkage of sexually antagonistic variants (reviewed in (Flintham and Mullon 2026). *P. angustifolia* sex chromosomes exhibit rearrangements despite distinguishing only males from hermaphrodites, indicating that the process can proceed even in the absence of full separation between sexes, hence in a situation where sexual antagonism may be expected to be weak. Sequencing additional individuals will be necessary to establish generality of the strikingly contrasted levels of rearrangements of the Y chromosomes we sequenced from *P. angustifolia* and *P. latifolia*. The oldest strata of the *P. angustifolia* Y chromosome (strata B2) seems to have become established at about the same time as the separation between the two species, so it is possible that the rearrangement process started to develop in one species but not the other. It is possible that their different life histories, *P. angustifolia* being a shrub and *P. latifolia* a tree, may play a role in this markedly different configuration. It is also possible that sexual antagonism (although inconspicuous at the phenotypic level) might be stronger in *P. angustifolia* than in *P. latifolia*. Alternatively, recent models have shown that sex chromosomes accumulate large-scale chromosomal rearrangements due to the fixation of rare “lucky” inversions carrying a less-than-average load of deleterious recessive mutations. The greater prevalence in one species as compared to the other may thus also be a simple reflection of the stochasticity of this evolutionary process (Lenormand and Roze 2022).

## Methods

### Plant material and sex phenotyping

Sex phenotypes of the samples, due to the evident morphological differences between hermaphrodite and male flowers (Fig. 1A), was determined by eye and confirmed under binoculars. For this work, we collected mixed-stage flower buds from *P. angustifolia* individuals before flowering for expression analysis. These individuals originate from a local population in Fabregues (southern France). We collected plant material from a *P. latifolia* male from the CEFE experimental garden in Montpellier (southern France) plant for genome assembly. We also collected leaves from *P. angustifolia* and *P. latifolia* individuals in several locations of France and Spain (see Table 1) during flowering season and observed the flower morphology of several branches to assign a sexual phenotype for each individual. Flower pictures were taken with the Keyence digital microscope VHX-X1.

**Table 1.** Origin and composition of the sampled individuals of *P. latifolia* and *P. angustifolia*.

| Population | Location | Observed phenotype | Species | Number of individuals | Use |
|---|---|---|---|---|---|
| CEFE-CNRS | 43.64N, 3.86E | Male M3 | *P. latifolia* | 1 | Genome assembly |
|  |  | Male M1, M2 | *P. latifolia* | 2 | Transmission distortion |
| Cadiz (Spain) 2015 | 36.53N, 6.21W | Hermaprodite | *P. angustifolia* | 10 | KASPar genotyping |
|  |  | Male | *P. angustifolia* | 16 | KASPar genotyping |
| Roussières (France) | 43.75N, 3.71E | Hermaprodite | *P. latifolia* | 9 | KASPar genotyping |
|  |  | Male | *P. latifolia* | 10 | KASPar genotyping |
| Sainte Lucie (France) | 43.05N, 3.05E | Hermaphrodite | *P. angustifolia* | 9 | KASPar genotyping |
|  |  | Male | *P. angustifolia* | 11 | KASPar genotyping |
| Fabrègues (Fance) | 43.53N, 3.75E | Hermaprodite | *P. angustifolia* | 10 | illumina WGS |
|  |  | Male | *P. angustifolia* | 10 | Illumina WGS |

### Genotyping of natural populations of *P. angustifolia* and *P. latifolia*

We used the KASPar assay system to genotype wild Phillyrea individuals based on their known sex phenotype and verify the presence of the Y-SDR allele in the male individuals and its absence in the hermaphrodite individuals. Specifically we sampled 10 *P. angustifolia* hermaphrodites and 16 males from a population located in Cadiz (Spain) and 9 hermaphrodites and 11 males from a population located in Sainte Lucie (France). For *P. latifolia,* 9 hermaphrodites and 10 males were sampled from a population located in Roussières (France). The ANDY2-specific T-probe labelled with FAM™ dye generates a fluorescence signal detected in the 523-568nm window and the UIF1-specific C-probe labelled with HEX™ dye generates a fluorescence signal detected in the 483-533nm window using a LightCycler® 480 (Roche Diagnostics GmbH). Automatic allele calls were performed with the LightCycler 480 SW1.5.0 SP3 software version 1.5.0 (Fig. S13 and Table S7).

### DNA extraction of leaves

DNA was extracted from lyophilised leaves using NucleoMag Plant kit (Macherey Nagel, Germany), following the manufactury protocol that we improved and using an automated method of extraction: Kingfisher Flex (Thermofisher, USA). Briefly, 50mg of lyophilised leaves material are grounded with the MM400 Retsch. The leaves are lysed for 1 hour at 56°C in MC1 lysis buffer supplemented with 10µl of liquid proteinase K (Macherey-Nagel, Germany), followed by a centrifugation step. The DNA is then bound to the magnetic beads. After magnetic separation and removal of supernatant, the magnetic beads are washed and DNA is eluted in a low salt buffer. DNA quantities were measured using a Qubit (Thermofisher, USA).

### Illumina library preparation

The illumina libraries were constructed using the NEXTFLEX Rapid XP V2 DNA-Seq kit (Revvity, USA) according to manufactured recommendations. In the first step, 100ng of total DNA is fragmented using an enzyme cocktail for 5 minutes at 35°C, followed by end-repair and adenylation steps. Next, the DNA is ligated with universal adaptator P5/P7 containing a unique dual index (NEXTFLEX Unique Dual Index Barcodes, Revvity, USA), followed by a post-ligation magnetic bead cleanup step. The DNA is then amplified via PCR (6 cycles) and purified again using magnetic beads. The libraries are subsequently quantified using Qubit (Thermofisher, USA) and quality-checked via gel electrophoresis (DNF473, Agilent, USA) on the 5200 Fragmenter Analyzer (Agilent, USA). Finally an equimolar pool of the libraries is prepared at 10nM and sent to an illumina sequencing platform Novaseq X plus25B (Illumina, USA) (2×150pb, paired end).

### High Molecular Weight DNA isolation of *P. latifolia* male individual leaves

High molecular weight (HMW) DNA was extracted from frozen leaves using QIAGEN Genomic-tips 500/G kit (Qiagen, MD, USA), following the tissue protocol extraction. Briefly, 2g of young leaves material were grounded in liquid nitrogen with mortar and pestle. After 3h of lysis and one centrifugation step, the DNA was immobilized on a column. After several washing steps, DNA was eluted from the column, then desalted and concentrated by isopropyl alcohol precipitation. A final wash in 70% ethanol was performed and the DNA was resuspended in EB buffer. DNA quantity and quality were assessed using NanoDrop and Qubit (Thermo Fisher Scientific, MA, USA). DNA integrity was also assessed using the Agilent FP-1002 Genomic DNA 165 kb on the Femto Pulse system (Agilent, CA, USA).

### PACBIO sequencing and assembly of *P. latifolia* male genome

To construct the genome assembly of *P. latifolia,* high Fidelity (HiFi) libraries were constructed using the SMRTbellÒ Template Prep kit 2.0 (Pacific Biosciences, Menlo Park, CA, USA) according to PacBio recommendations (SMRTbellÒ express template prep kit 2.0 - PN: 100-938-900). HMW DNA samples were first purified with 13 Agencourt AMPure XP beads (Beckman Coulter, Inc, CA USA), and sheared with Megaruptor 3 (Diagenode, Liège, BELGIUM) at an average size of 20 kb. After end repair, A-tailing and ligation of SMRTbell adapter, the library was size-selected on the BluePippin System (Sage Science, MA,USA) at range sizes of 10-50kb. The size and concentration of libraries were assessed using the Agilent FP-1002 Genomic DNA 165 kb on the Femto Pulse system and the Qubit dsDNA HS reagents Assay kit. Sequencing primer v5 and SequelÒ II DNA Polymerase 2.2 were annealed and bound, respectively, to the SMRTbell libraries. Each library was loaded on two SMRTcell 8M at an on-plate concentration of 90pM. Sequencing was performed on the Sequel II system at the Gentyane Genomic Platform (INRAE Clermont-Ferrand, France) with Sequel II Sequencing kit 3.0, a run movie time of 30 hours with an Adaptive Loading target (P1 + P2) at 0.75. HiFi reads were produced with the PacBio Sequel II system on four SMRTCells and were assembled using the HiFiasm assembler (v0.15.5) (Cheng et al. 2021). To assess the completeness and quality of the assembly, we used the Benchmarking Universal Single-Copy Orthologs (BUSCO) pipeline with the viridiplantae database (Simão et al. 2015). We obtained a 99.3% complete BUSCO score.. In addition, we performed k-mer analysis to quality control the dataset using Jellyfish tool (Marçais and Kingsford 2011) and the assemblies using module ‘‘comp’’ of the k-mer Analysis Toolkit (Mapleson et al. 2017). Metrics are reported in the Table S3.

### RNA-seq library construction and sequencing

The TruSeq RNA sample Preparation v2 kit (Illumina Inc., USA) was used according to the manufacturer’s protocol with the following modifications. In brief, poly-A containing mRNA molecules were purified from 1 ug total RNA using poly-T oligo attached magnetic beads. The purified mRNA was fragmented by addition of the fragmentation buffer and was heated at 94 ॰C in a thermocycler for four minutes to yield library fragments of 250-500 bp. First-strand cDNA was synthesized using random primers to eliminate the general bias towards 3’end of the transcripts. Second strand cDNA synthesis, end repair, A-tailing, and adapter ligation was done in accordance with the manufacturer’s protocols. Purified cDNA templates were enriched by 15 cycles of PCR for 10s at 98 ॰C, 30s at 65 ॰C and 30s at 72 ॰C using PE1.0 and PE2.0 primers and with Phusion DNA polymerase (NEB, USA). Each indexed cDNA library was verified and quantified using a DNA 100 Chip on a Bioanalyzer 2100, then pooled in equimolar amounts. The final library was quantified by real-time PCR with the KAPA Library Quantification Kit for Illumina Sequencing Platforms (Kapa Biosystems Ltd, SA) adjusted to 10 nM in water and provided to the Get-PlaGe core facility (GenoToul platform, INRA, Toulouse, France http://www.genotoul.fr) for sequencing. Final pooled cDNA libraries were sequenced using the Illumina mRNA-Seq, paired-end protocol on a HiSeq2000 sequencer, for 2 x 100 cycles. Libraries were diluted to 2 nM with NaOH and 2.5 mL transferred into 497.5 mL HT1 to give a final concentration of 10 pM. 120 mL were then transferred into a 200 mL strip tube and placed on ice before loading onto the cBot. Mixed libraries from ten individual indexed libraries were run on a single lane. The flow cell was clustered using TruSeq PE Cluster Kit v3, following the Illumina PE_Amp_Lin_Block_V8.0 protocol. Following the clustering procedure, the flow cell was loaded onto the Illumina HiSeq 2000 instrument. The sequencing chemistry used was v3 (FC-401-3001, TruSeq SBS Kit) with 2×100 cycles, paired-end, indexed protocol.

### Annotation of protein-coding genes for haplotype resolved assemblies

We conducted a liftoff (Shumate and Salzberg 2021) of the primary annotation of *P. angustifolia* previously published (Castric et al. 2024) to annotate the genes of the two haplotype-resolved assemblies used in this study: 040A11_h1 and 040A11_h2 for the male assembly (040A11) and 01N25_h1 01N25_h2 for the hermaphrodite assembly (01N25). We were able to lift over most of the gene models (Table 2) from the initial of 25,913 from the original annotation. Next, we obtained a set of proteins of the Olive tree (*Olea europaea*) from (Julca et al. 2020) (OE9D.longest_peptide.fa; 55,732 proteins) to complement the liftover using miniprot (Li 2023). First, we masked the genome using the liftoff gene model positions to not reannotate those, then, after miniprot ran, we retained the longest hit per loci and discarding mRNAs with in-frame stop codons at CDS with GFFread (Pertea and Pertea 2020). This approach added around ∼16,000 more gene models to the assemblies (see Table 2). Finally, we aimed to functionally annotate this set of genes using two parallel strategies. First, by searching for homologs from Araport11 (Cheng et al. 2017) and assigning each gene model its closest Araport11 homolog. For genes annotated via liftoff, we additionally included the corresponding OLEA9 homologs from *Olea europaea* database (Julca et al. 2020). Second, we performed domain-based annotation using InterProScan to further characterize the predicted gene functions. We ran InterProScan locally using the following databases: CDD, PANTHER, Pfam, SUPERFAMILY and TIGRFAM and we were able to assign some level of function to around 86% of all annotated genes (see Table 2 for details).

**Table 2.** Summary statistics of the gene and functional annotation of the haplotype resolved genomes.

|  | 01N25 hap1 | 01N25 hap2 | 040A11 hap1 | 040A11 hap2 |
|---|---|---|---|---|
| Liftoff | 24,942 | 24,900 | 25,709 | 25,637 |
| Miniprot Olea9 | 15,881 | 15,838 | 16,027 | 15,853 |
| Total annotated | 40,823 | 40,738 | 41,736 | 41,490 |
| interproScan | 35,478 | 35,112 | 36,384 | 35,889 |
| Araport11 homologs | 33,649 | 33,373 | 34,233 | 33,811 |
| OLEA9 homologs | 24,040 | 23,829 | 24,721 | 24,378 |

### Annotation of repetitive elements for primary and haplotype resolved assemblies

To annotate transposable elements in *Phillyrea angustifolia*, we opted for the de novo approach of using EDTA (Ou et al. 2019) on the primary assemblies of both the male (040A11) and the Hermaphrodite (01N25) individuals. We ran EDTA with the options “*-species others --step all --sensitive 1 --force 1 -anno 1 --evaluate 1*” to obtain a separate TE library for each assembly. Then, we ran TRASH (Wlodzimierz et al. 2023) and TRF (Benson 1999) on the primary assemblies of both individuals to obtain the annotation of centromeres, TRASH, and other satellite repeats, TRF, using default settings. By converting the output of both tools to GFF and merging them, we obtained the satellite annotation for the primary assemblies. We applied the same method to obtain the satellite annotation of the haplotype resolved assemblies. We also improved the quality of the TE annotation by running TEtrimmer (Qian et al. 2024). First, using the satellite annotation, we masked the primary assemblies of both individuals merging annotated features up to 1 kb apart using with bedtools ‘bedtools merged 1000’ with the aim of masking the centromeres and simple repeats of the assemblies to enhance the quality of the automatic TE curation. Next, we used seqkit (Shen et al. 2024) to merge both TE libraries of the primary assemblies and remove duplicated TE models. Finally, we ran TEtrimmer on the merged TE library and on a merged,masked assembly of both individuals. This final TE library product of TEtrimmer was manually inspected with TE-AID (Goubert et al. 2022) to remove spurious TE models with huge increases in size (more than 15 kb) after the TEtrimmer run. We finalized the TE annotation by removing TE models categorized as Unknown by TEtrimmer. Using the newly constructed TE library, we annotated the haplotype resolved assemblies of 040A11 and 01N25 using repeatmasker (Smit, AFA, Hubley, R & Green, P. 2013) with the options: “-*e ncbi -s -a -inv -xsmall -div 40 -no_is -nolow -cutoff 225 -norna -gff*”, we polished the output of repeatmasker by passing the tool One code to find them all (Bailly-Bechet et al. 2014) with the option “*--unknown*” and finally, we manually removed TEs that overlapped with their corresponding satellite annotation and also those that were smaller than 80bp. Finally, we wanted to screen for full length LTR retrotransposons in the *P. angustifolia* haplotype resolved assemblies. For this, we used LTRpred (Drost 2020) using the tRNAs, Dfam and HMM models provided for LTRpred and the mutation rate of 7.7e^-09^ used for the olive tree (Olea europaea) (Julca et al. 2020). We masked the assemblies using their corresponding satellite annotation before running LTRpred to avoid false detection of LTRs.

### Sex determining region discovery and annotation

We used minimap2 (Li 2018) with the preset “asm5” to find structural variants across the different haplotypes from the hermaphrodite and the male haplotype resolved assemblies published in (Castric et al. 2024), Next, aim to associate the rearrangements found at chromosome 12 with the sex phenotype. First, we remapped the previously published sex-linked markers from (Carré et al. 2021) with *bwa mem* (Li and Durbin 2009) using default settings and verified the location of these markers being within and in close proximity to the heavily rearranged region at chromosome 12. To further validate this region as the SDR of P. angustifolia, we remapped the previously published RAD-seq raw data (Carré et al. 2021) using a combination of Stacks (Catchen et al. 2013) and bwa-mem (Li and Durbin 2009). We made use of the function process_radtags of Stacks (Catchen et al. 2013) with the options “*--renz-1 ‘pstI’ --renz-2 ‘mseI’ -E phred33*” to process the RAD-seq paired data and then, using bwa-mem program (Li and Durbin 2009) with default parameters, we mapped these reads against the male haplotype resolved assemblies (040A11.h1 and 040A11.h2). Finally, we calculated the read coverage (10kb windows) using mosdepth (https://github.com/brentp/mosdepth) for each sample of these two references. For comparison, we separated male and hermaphrodite samples in two groups and calculated the mean coverage of each of the groups for each window. We visualised these results by log2 transforming the average coverage of each window and applying the function geom_smooth “method ‘gam’; formula = ‘y ∼ s(x, bs = “cs”)” from the ggplot2 package. To annotate the SDR region in chromosome 12 predicted by the structural variants and the cosegregation analysis we used GENESPACE (Lovell et al. 2022) with default parameters. GENESPACE uses MCScanX to group protein similarity scores into syntenic blocks and employs Orthofinder (v2.5.4) to identify haplotype homologs within these synteny-constrained regions. In our case, we used GENESPACE to search for disruption of synteny using the proteins of the four haplotypes of the two *P. angustifolia* assemblies with a special focus on the structural variant located at chromosome 12. To identify gene relationships between Y- and X-SDR alleles. We first defined private Y- and X-linked genes as genes located in the SDR that lack a homolog in the corresponding SDR region of the opposite allele. Because GENESPACE can be confounded by gene duplication events, we extracted protein sequences at the SDR for both alleles (146 genes for Y-allele and 117 genes for X-allele and ran a reciprocal diamond blastp with the options: “-*-ultra-sensitive” and “--max-target-seqs 1*”, We defined as homologs gene pairs which reciprocal hit has a percentage of identity of more than 80%, below that threshold, including no matches, we define those genes as private for each allele. We also define homolog gene pairs as syntenic or rearranged if both members of the pair belong to the same syntenic block as defined by GENESPACE, otherwise we classify those homologs as rearranged.

### Validation of the SDR in a natural population and evaluation of its genetic diversity

We collected leaf samples from 21 individuals collected in the form of seeds produced through open pollination within a natural population of *P. angustifolia* in Fabregues (France; 43°28’34“N; 3°45’37”E) and hosted on the “Plateforme des Terrains d’Expérience du LabEx CeMEB” (CEFE, CNRS). Paired-end Illumina raw reads were trimmed and filtered using Trim Galore (Krueger 2015), and quality control was assessed with FastQC (Andrews et al. 2012). We then used Meryl (Rhie et al. 2020) to extract 21-mers from the reads of each individual sample. Samples were subsequently grouped according to their sex phenotype (males and hermaphrodites), and a k-mer–based discriminatory analysis was performed. Specifically, we identified 21-mers that were present in all individuals of one phenotype and absent from all individuals of the other phenotype. This analysis yielded 152 hermaphrodite-specific 21-mers and 1,146,472 male-specific 21-mers. Next, clean reads were aligned to the male reference genome (040A11), considering haplotype 1 (040A11.h1) and haplotype 2 (040A11.h2) separately, using BWA-MEM (Li and Durbin 2009) with ‘*-k 21 -T 21 -a -c 1000*’ parameters. 1146134 21-mers were mapped to the genome, and 338 were not (99% mapped). Finally, we summarized these results by computing the number of mapped k-mers per 100 bp windows for analysis and 100 Kbp windows for visualization purposes using bedtools (Quinlan and Hall 2010). Variant calling was then performed independently for each haplotype using the clean reads. We used bcftools mpileup (Danecek and McCarthy 2017) with the parameters ‘*--redo-BAQ --min-BQ 20 -d 200*’ followed by variant calling with *‘bcftools call -c*’. Variants were subsequently normalized and filtered using a combination of bcftools and VCFtools (Danecek et al. 2011). Specifically, we applied‘*bcftools norm -m -any -c x*’ and *‘vcftools –gzvcf --maf 0.05 --max-missing 0.2*’, while retaining invariant sites. Finally, we obtained population genetic summary statistics and their distribution across the genome with pixy (Korunes and Samuk 2021) using a window size of 50 Kbp.

### dN/dS analysis

Using single copy syntenic gene pairs (orthologs) identified by GENESPACE in both haplotypes of the male reference (040A11) we investigated patterns of allelic divergence between both haplotypes. For each pair, we used the Smith-Waterman algorithm to align both proteins and then revtrans (Wernersson and Pedersen 2003) to extract the codon-wise aligned CDS. We then used Kaks_calculator 3.0 (Zhang 2022) using the codon-aware method of Goldman and Yang (Goldman and Yang 1994) to estimate the ratio of neutral substitutions per site (dS). Because the number of dS estimates differed greatly between SDR alleles (55 values) and the genomic background (31,579 values), we tested whether the observed difference in medians could be explained by sampling size alone. To do so, we randomly sampled 55 values from the genomic background 10,000 times, calculated the median for each sample, and compared these bootstrapped medians to the observed median at the SDR using a Wilcoxon test. All 10,000 tests yielded significant results (p < 0.05). Figure S6 shows the distribution of bootstrapped medians relative to the observed SDR median. We repeated the same test using the dS values obtained when measuring the divergence between orthologs of *P. angustifolia* and *P. latifolia* and compared these with the SDR median values of *P, angustifolia*.

### *P. latifolia* SDR characterization and cross species comparison

To make comparisons between chromosome 12 haplotypes of the male assembly of *P. angustifolia* (040A11) and the assembly of *P. latifolia* (also a male individual). We used minimap2 (Li 2018) with the preset option “-*asm5*” to align the four haplotypes and then used pafR (https://github.com/dwinter/pafr) to visualize the alignments and to retain only the primary alignments with a mapping quality of 60 and convert them to bed files. Then, we used bedtools (Quinlan and Hall 2010) to extract the boundaries of the SDR alleles in *P. latifolia* X and Y SDR alleles by cross comparison. Once defined, we used these boundaries to extract the TE and gene information from the annotation at these loci. Lastly, we did a cross-species comparison of both Y and X homologous alleles to extract common genes between both species at the SDR and compared them. We use diamond (Buchfink et al. 2015) to perform protein to protein alignments of genes of both *P. latifolia* SDR alleles against the *P. angustifolia* genes from both SDR alleles. We filtered matches by applying a filter of 80% protein identity over 80% of subject length.

### Gene expression analysis

We sampled flower buds from 14 hermaphrodite and 13 male *P. angustifolia* individuals to examine patterns of gene expression during three flower buds representing three stages 3, 1, 0 days respectively before flower opening. Raw paired-end FASTQ reads were trimmed and filtered with Cutadapt (Martin 2011) using default parameters. To limit reference bias, we concatenated both haplotypes from the male assembly (040A11), containing the 2 alleles of the SDR, into a single reference and mapped the processed paired-end reads onto it using HISAT2 (Kim et al. 2019) with default parameters. Gene expression levels were quantified with th*e htseq-count* function from HTSeq (Anders et al. 2015; Kim et al. 2019) with the option “*--nonunique none*“ and applied separately to the gene annotations of haplotype 1 and haplotype 2. The resulting per-sample counts were combined into a matrix using a custom Python script independently for each haplotype. Three samples were filtered as they show a high number of not aligned reads or total reads and another one due to mislabeling. Additionally, we pre-filtered low count genes with less than 10 counts in at least 10 samples as suggested (Love et al. 2014) by the DESeq2 manual. Finally, differential expression analyses comparing hermaphrodite and male samples were then performed independently for each haplotype using DESeq2 (Love et al. 2014). We considered differentially expressed genes those with an adjusted p-value of less than 0.01 between the two groups (Benjamin and Hochberg method) and a log2 fold change above 1.

### Gene model extension

Upon visual inspection of the annotation tracks, we realized the gene model for the gene “maker-chr_M_12-augustus-gene 106.15” was not consistent with the pattern of RNA-seq reads mapping. We manually fixed this gene model by extracting the reads of the gene region (040A11.chr12.h1-12:10682116-10684970) only for the Ha individuals (since those individuals show high levels of expression for that gene). Then, we used Stringtie to create a new gene model from these Ha transcripts, we then manually extended the borders to capture the entire CDS and set the correct phase. We used *agat_sp_fix_cds_phases.pl* script from the AGAT suite to generate a full GFF record for that gene model and replace it in the annotation GFF file. Finally, we used miniprot to liftover this new gene model to the rest of the assemblies.

### Phylogenetic analysis of gene families

To gain insight into the evolutionary origins of ANDY1/2 and the Ole e1 gene families, we conducted a phylogenetic reconstruction of these two gene families across Oleaceae. We used publicly available proteomes of seven species spanning major clades of Oleaceae: *Forsythia ovata, F. suspensa, Abelliophyllum distichum, Syringa oblata, Fraxinus pennsylvanica, Fraxinus excelsior, Olea europaea* and a species from the sister genus of Phillyrea, *Osmanthus fragrans* (Dupin et al. 2024), which is a known androdioecious species. We completed this dataset incorporating the proteomes predicted from the two haplotypes of both *P. angustifolia* and *P. latifolia*. Identification of gene families and the phylogenetic inference into groups of orthologs and paralogs was done using Orthofinder (Emms and Kelly 2019). For each protein family, we retrieved the corresponding orthogroup IDs for the different species and extracted both the CDS and the proteins. We aligned the protein sequences obtained using MAFFT (Katoh and Standley 2013) with options ‘--maxiterate 1000 --localpair’. We then converted these amino acid alignments back into codon alignment using RevTrans (Wernersson and Pedersen 2003) and their corresponding CDS sequences. We trimmed the alignments using trimAl (Wernersson and Pedersen 2003; Capella-Gutiérrez et al. 2009) with the option ‘-gt 0.8‘. We manually curated the Ole e1 alignment using MACSE v2.00 (Ranwez et al. 2011) during which we removed 16 putative orthologs from *S. oblata* that were erroneously identified as orthologs on the Ole e1 gene family. Finally, we constructed a maximum likelihood tree of each gene family with IQ-tree 3 (Wong et al. 2025) with parameters ‘-m TEST -B 1000 -T AUTO -st CODON. The model MG+F3X4+G4 was selected for the ortholog family of Ole e1, reporting a BIC score of 14433.672, Whereas the model of the UIF1 ortholog gene family was MG+F1X4+G4, with a BIC score of 8604.534. Due to the high similarity between sequences, high bootstrap node support was difficult to obtain for the majority of the nodes of the Ole e1 gene family, thus we desisted to date the duplication events of these paralogs. For the UIF1 gene family, gene duplication events were identified using MEGA’s Find Gene Duplications function (Zmasek and Eddy 2001), which distinguished duplication events from speciation events in the gene tree. To estimate the duplication ages, MEGA’s RelTime-ML analysis (Tamura et al. 2018) was performed on the gene tree, using the known species divergence times to calibrate the corresponding internal speciation nodes as previously reported for the Oleaceae family (Dupin et al. 2022; Dupin et al. 2024). All speciation calibration nodes were modeled under a normal distribution with a standard deviation adjusted to comprise the 95% CI of the calibration dates. The mean values (in millions of years) were as follows: speciation between *P. angustifolia* and *P. latifolia* set to 8.5 MYA; divergence of Phillyrea and Osmanthus genus set as 17.5; diversification of the Oleinae set as 33.7; divergence between Fraxininae and Oleinae set as 48.5; diversification of the Ligustrinae from Fraxininae and Oleinae set as 53.0; diversification of Forsythieae from the rest of genus set as 78.0. Confidence intervals are displayed in Fig S14. Finally, phylogenetic trees were visualised with the R package ggtree (Yu et al. 2017).

## Data and resource availability

The data generated during this study are publicly available in the following repositories: Raw short read genomic datasets have been deposited in XXXXX. PacBio Hifi reads and assembly of P. latifolia are deposited in XXXXX. Expression data is deposited in XXXXX. All scripts used in this study are publicly available on the following link: https://gitlab.univ-lille.fr/eep_lab/vincent_castric/Androdiecy_SDR.

## Acknowledgments

The authors thank the Hauts-de-France Region, the Ministry of Higher Education and Research, and the European Fund for Regional Economic Development for their financial support to the CPER ECRIN program. We also acknowledge funding from the European Research Council (NOVEL project, grant #648321) and the Agence Nationale de la Recherche (project TE-MoMa, grant ANR-18-CE02-0020-01). The French State under the France-2030 programme and the Initiative of Excellence of the University of Lille are acknowledged for the funding and support granted to the R-CDP-24-002-PIE project. We thank Thomas Lesaffre for comments on the manuscript.

## Author Contributions

Conceptualization, A.C.G., P.S.-L., P.V., V.C. and S.B.; methodology, A.C.G., C.G., C.M., P.S-L and C.P; investigation, A.C.G, C.P, M.M A.V., A.B., and J.W. resources: P.S.-L., M.D., V.C.; writing original draft: A.C.G. V.C. and P.S.-L.; writing – review & editing, A.C.G., V.C., P.S.-L., S.B., X.V., P.V., J.W., M.D.; funding acquisition, P.S.-L. and V.C.; supervision, V.C.

