## Supplementary_material for "The structure and evolution of sex chromosomes in an androdioecious species with two mating types"

### Supplementary Tables

**Table S1.** Summary statistics of the new annotation of the *P. angustifolia* assemblies published in (Castric et al. 2024).

| | 01N25 hap1 | 01N25 hap2 | 040A11 hap1 | 040A11 hap2 |
|---|---|---|---|---|
| Number of genes | 40,823 | 40,736 | 40,699 | 40,521 |
| % Satellite | 2,1% | 2.6% | 1.9% | 2.5% |
| Number of Satellite | 139,508 | 165,696 | 141,577 | 163,954 |
| % TE | 55.6% | 55.0% | 55.6% | 54.9% |
| Number of TEs | 607,603 | 604,673 | 609,860 | 597,246 |
| Intact LTR sampled | 14,928 | 14,805 | 14,904 | 14,351 |
| Total assembly size (Mbp) | 773.38 | 771.75 | 773.05 | 759.85 |
| Total SDR size (Mbp) | 5.25 | 5.48 | 6.83 | 5.31 |

**Table S2**. Sex ratio in the progeny of crosses between two *P. angustifolia* hermaphrodites (13A30 and 01N04) and two *P. latifolia* males (M1 and M2). Sex of the offspring (embryos) was determined by the KASPar marker tracking the presence of the ANDY1/2 genes. H indicates hermaphrodites and M males. The likelihood of Mendelian segregation of the sex phenotype was tested using the binom.test function in base R, assuming an expected 1:1 segregation ratio (50% probability for each sex phenotype).

| Pollen recipient (*P. angustifolia* hermaphrodite) | Pollen donor (*P. latifolia* male) | Offspring | Probability of deviation from mendelian segregation |
|---|---|---|---|
| 13A30 [Hb] | M1 | 0 H / 56 M | $2.78e^{-17}$ |
| 01N04 [Ha] | M1 | 5 H / 1 M | 0.219 |
| 13A30 [Hb] | M2 | 1 H / 37 M | $2.84e^{-10}$ |
| 01N04 [Ha] | M2 | 12 H / 6 M | 0.238 |

**Table S3.** Assembly and annotation statistics of the male *P. latifolia* genome.

| | Haplotype 1 | Haplotype 2 |
|---|---|---|
| | Assembly statistics | |
| Contig number | 35 | 45 |
| Megabase level contigs | 27 | 36 |
| Total size (bp) | 750,934,917 | 736,987,348 |
| Largest contig (bp) | 52,522,564 | 43,341,961 |
| GC content % | 34.52 | 43.45 |
| N50 | 31,438,971 | 30,816,052 |
| | Liftoff statistics | |
| Gene number | 45,904 | 45,776 |
| Average coding sequence length | 1060 | 1059 |
| Average exon number | 4.7 | 4.7 |
| | Functional annotation | |
| CDD | 13,544 | 13,559 |
| Panther | 37,704 | 37,643 |
| Pfam | 32,997 | 32,966 |
| Superfamily | 25,275 | 25,223 |
| Total number of annotated genes | 39,075 | 39,023 |

**Table S4.** Set of conserved genes at the Y-SDR allele in the Phillyrea genus and their expression status in mixed-stage male flower buds in *P. angustifolia*.

| ***P. latifolia* gene ID** | ***P. angustifolia* gene ID** | **Putative gene function** | **Average expression counts in *P. angustifolia* males** |
|---|---|---|---|
| H1_MP002491 | MP002491 | - | 0 |
| H1_MP005022 *and* H1_maker-chr_M_12-augustus-gene-171.1 | augustus_masked-chr_M_12-processed-gene-136.2 | Pollen allergen Ole e 1 family | 11831 |
| H1_MP007794 | MP007794 | - | 0 |
| H1_MP014556 | MP014556 | Calcium-permeable cation-selective channel. | 0 |
| H1_MP014741 | MP014741 | CCR4-NOT transcription complex subunit. | 0 |
| H1_MP017106 | MP017106 | - | 0 |
| H1_augustus_masked-chr_M_12-processed-gene-136.0 | augustus_masked-chr_M_12-processed-gene-136.0 | Pollen allergen Ole e 1 family | 6169 |
| H1_augustus_masked-chr_M_12-processed-gene-95.2 | augustus_masked-chr_M_12-processed-gene-95.2 | Myb family transcription factor HRS1-like | 273 |
| H1_maker-chr_M_12-augustus-gene-95.26 | maker-chr_M_12-augustus-gene-95.26 | Myb family transcription factor HRS1-like | 231 |

**Table S5**. Hypergeometric distribution test of enrichment of differentially expressed genes within the sex-determining region (SDR) of *P. angustifolia*.

| Haplotype | Total expressed genes | Total DEGs | SDR expressed genes | SDR DEGs | Hypergeometric test p-value |
|---|---|---|---|---|---|
| 040A11.h1-Y | 17,894 | 343 | 70 | 49 | $1.643677e^{-72}$ |
| 040A11.h2-X | 17,394 | 257 | 67 | 31 | $2.119079e^{-39}$ |

**Table S6**. Origin of the publicly available proteomes from Oleaceae used in this study.

| Species | Source publication |
|---|---|
| *Abelliophyllum distichum* | Chromosome-level assemblies of the endemic Korean species *Abeliophyllum distichum* and *Forsythia ovata*. (Jang et al. 2024) |
| *Syringa oblata* | A chromosome-level genome of Syringa oblata provides new insights into chromosome formation in Oleaceae and evolutionary history of lilacs. (Wang et al. 2022) |
| *Forsythia ovata* | Chromosome-level assemblies of the endemic Korean species *Abeliophyllum distichum* and *Forsythia ovata*. (Jang et al. 2024) |
| *Forsythia suspensa* | The gap-free genome of Forsythia suspensa illuminates the intricate landscape of centromeres. (Cui et al. 2024) |
| *Fraxinus pennsylvanica* | A high-quality reference genome for Fraxinus pennsylvanica for ash species restoration and research. (Huff et al. 2022) |
| *Fraxinus excelsior* | Genome sequence and genetic diversity of European ash trees. (Sollars et al. 2017) |
| *Olea europaea* | Genome of wild olive and the evolution of oil biosynthesis. (Unver et al. 2017) |
| *Osmanthus fragrans* | The chromosome-level quality genome provides insights into the evolution of the biosynthesis genes for aroma compounds of Osmanthus fragrans. (Yang et al. 2018) |

**Table S7**. Probe sequences used for the KASPar genotyping assay (probe 889).

| | |
|---|---|
| T-probe-FAM™ | ACAAGGCCCTCTGCRCCCT |
| C-probe-HEX™ | ACAAGGCCCTCTGCRCCCC |
| Common_reverse_primer | TATCATCATCTTCTTCCATGYTGTTTCGAT |

#### Supplementary figures

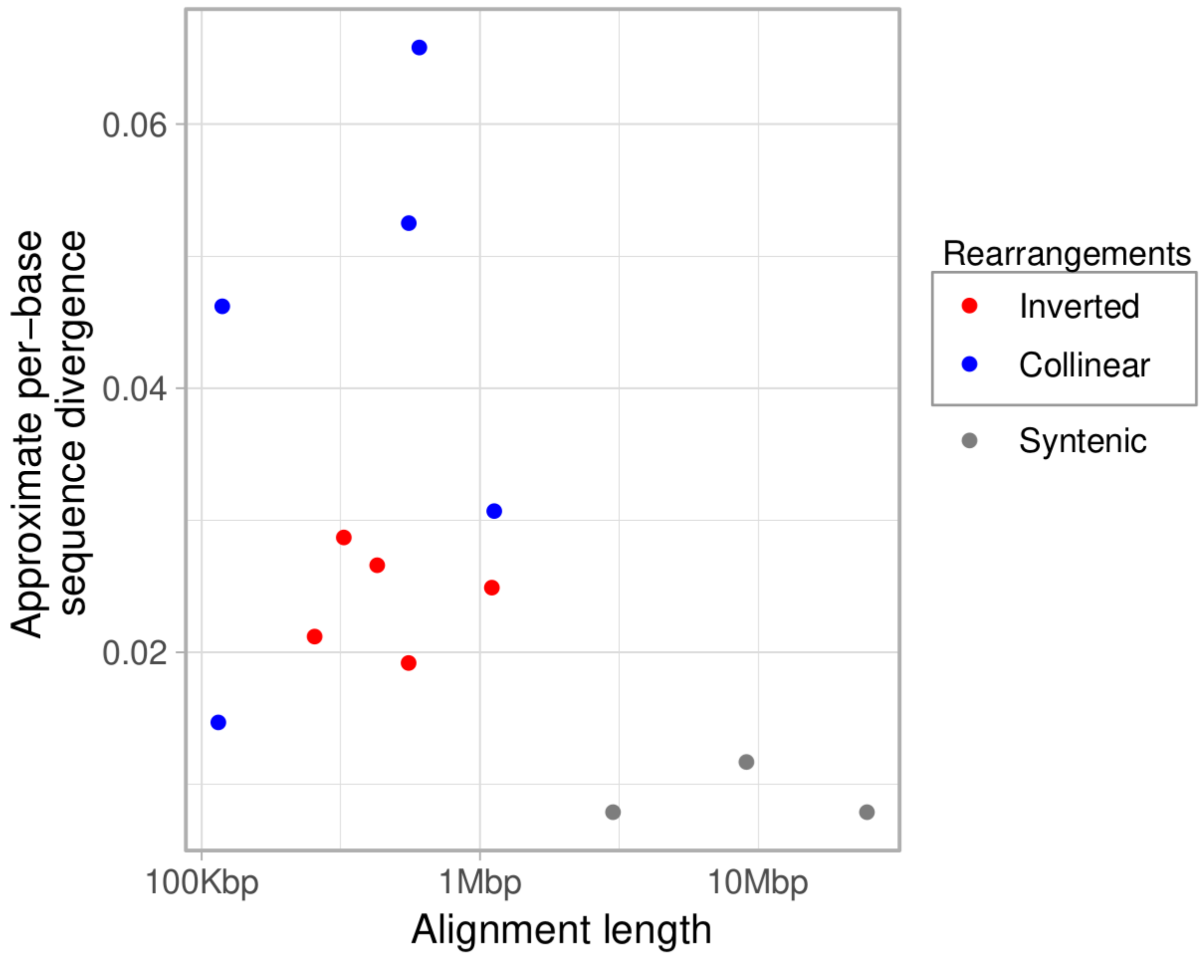


**Figure S1.** Alignment comparison of syntenic and SDR rearrangements between the X- and Y-containing haplotypes of *P. angustifolia* accession 040A11. These alignments were produced by minimap2 which also computes the approximate per-base sequence divergence by default.

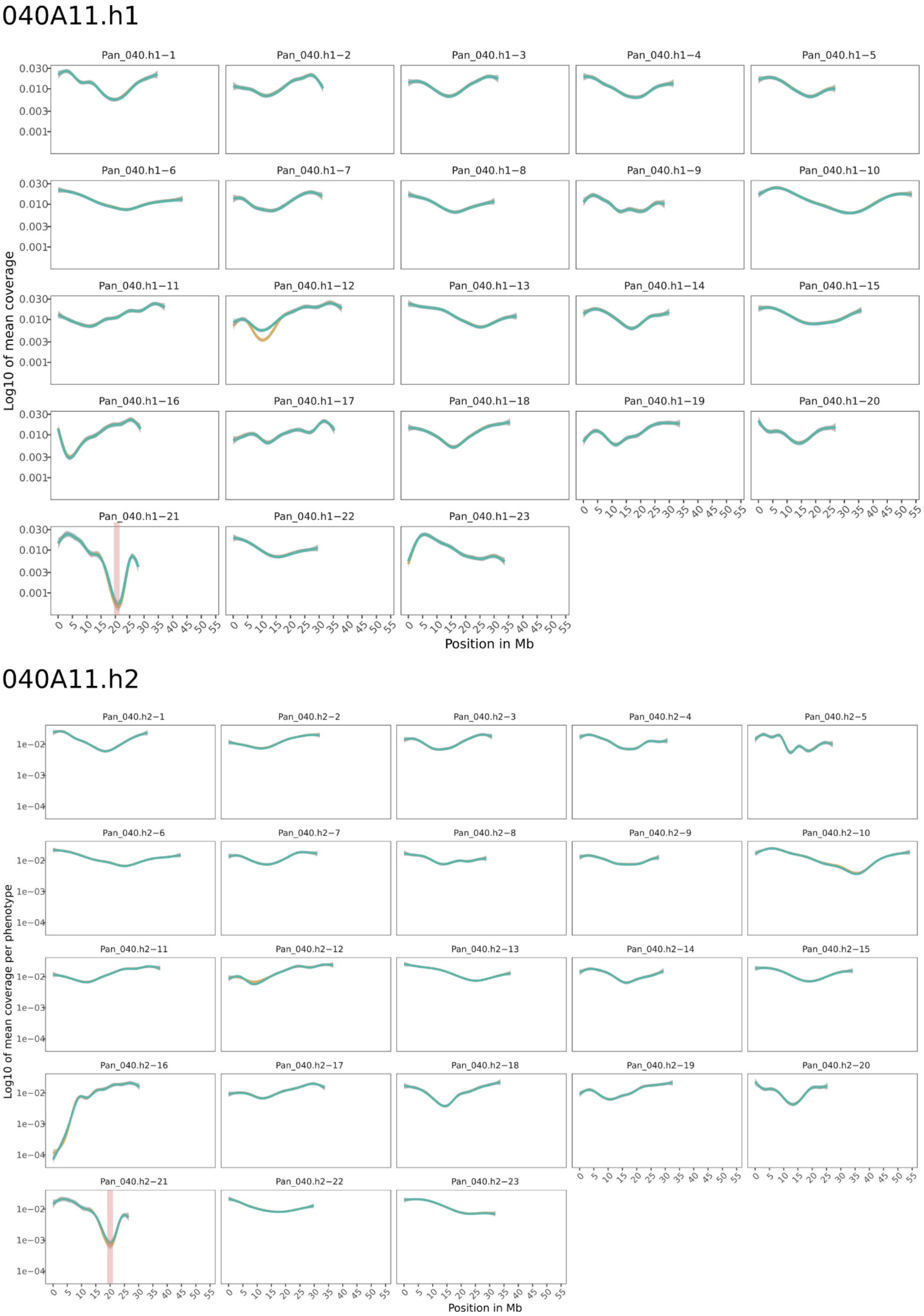


**Figure S2. Genome wide coverage across the 32 chromosomes.** Coverage comparison between hermaphrodites and males across the genome. The centromeric region in chromosome 21 is marked with a vertical red bar. Excluding the centromere, only high discrepancy was observed at chromosome 12 at the SDR.

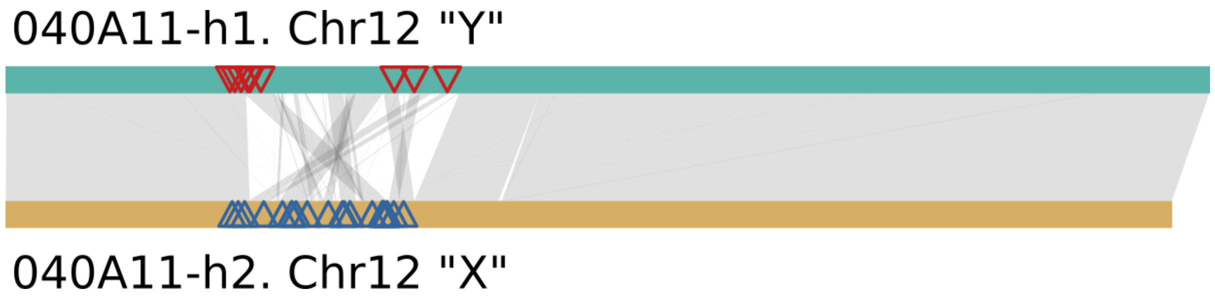


**Figure S3. Genomic placement of the associated sex markers.** Mapping of the published RAD-seq genetic markers associated with sex from (Carré et al. 2021) at both haplotypes of the male individual 040A11.

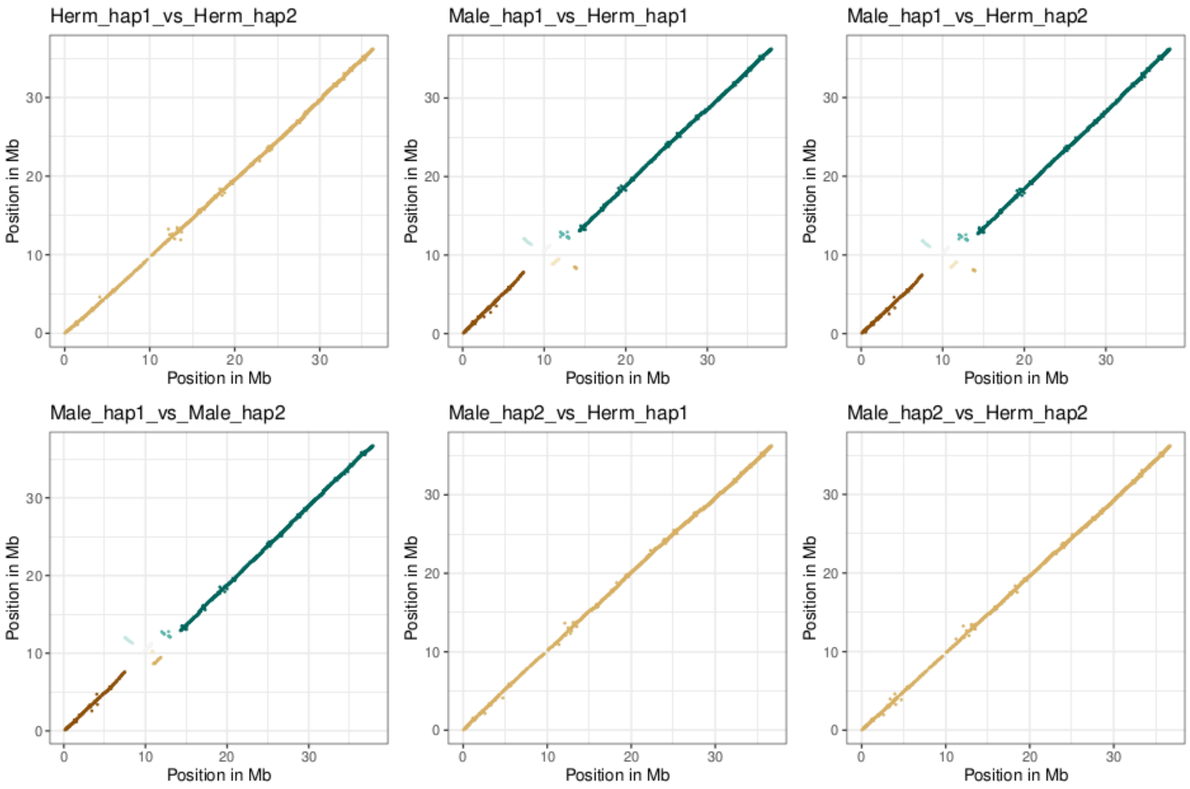


**Figure S4. Chromosomal syntenic relationships among the four haplotypes of the Male and Hermaphrodite assemblies of *P. angustifolia*.** Each panel shows pairwise syntenic blocks between corresponding haplotypes of chromosome 12. Points represent identified alleles between two genomes,displayed according to their positions (in megabases) along each haplotype. Colors denote distinct syntenic block IDs (blkID), as defined by GENESPACE. The diagonal patterns in red and purple represent total collinearity, while the rearrangements displayed in different colors represent translocation and inversions of the syntenic gene blocks between the “X” and “Y” haplotypes.

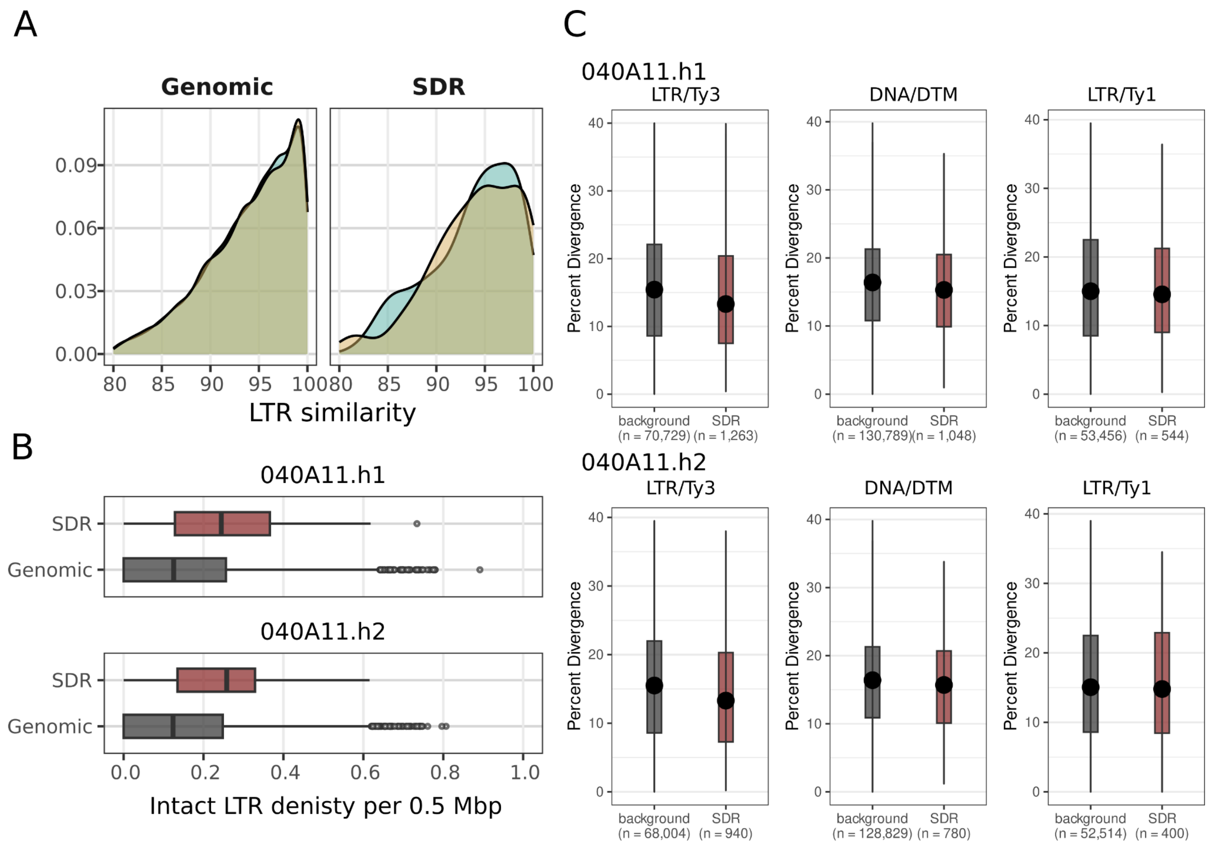


**Figure S5. *P. angustifolia* SDR harbours younger TEs than the genomic average for both alleles in 040A11. A)** Relative abundance of LTR age in both the SDR and the genomic background. Golden represents the haplotype with the "X" allele and turquoise the "Y" allele. **B)** Density of intact LTRs obtained using LTRpred at the SDR region and at the rest of the genomic background. **C)** Using the "divergence from consensus" metric as a proxy for TE age, we compared the age distributions of transposable elements (TEs) in the genomic background with those located at the SDR for the three largest superfamilies at the SDR: Mutators (DNA/DTM), Ty1 (LTR/Ty1), and Ty3 (LTR/Ty3). Using a Wilcoxon rank-sum test we estimated significant values at haplotype 1 for DTM (p-value $4.37e^{-08}$) and Ty3 (p-value $4.87e^{-07}$) but not for Ty1 (p-value 0.45); similar results were obtained for haplotype 2: DTM (p-value $2.26e^{-02}$) and Ty3 (p-value $6.99e^{-08}$), Ty1 (p-value 0.9).

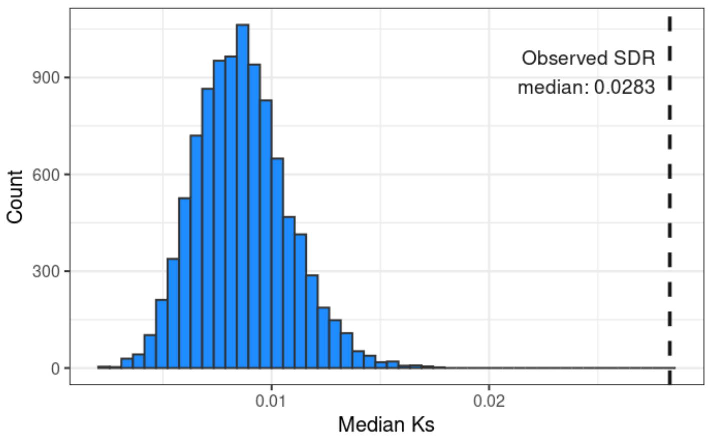


**Figure S6. The SDR alleles are more divergent from one another than genes in the genomic background.** The dashed line indicates the median dS observed between alleles of the 55 genes in syntenic blocks at the SDR. The histogram shows the distribution of median dS values obtained by randomly sampling 55 genomic background values 10,000 times.

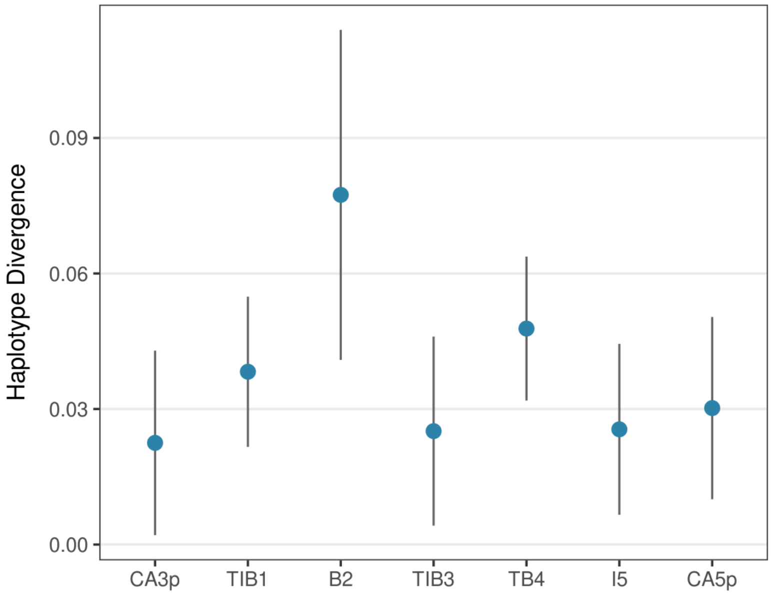


**Figure S7. DNA divergence between “Y” and “X” syntenic blocks at the SDR.** We used minimap2 to align by pairs the DNA sequences of the syntenic blocks of the “Y” and “X” alleles and averaged the “gap-compressed per-base sequence divergence” estimations given by minimap2.

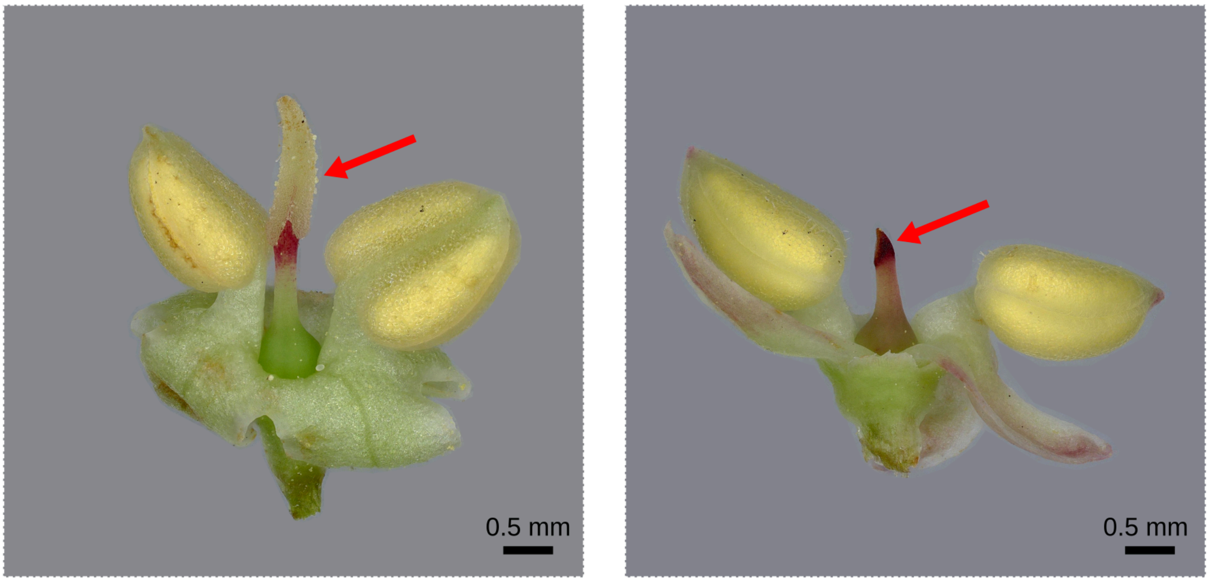


**Figure S8. Flower morphs of hermaphrodite and males of *P. latifolia*.** As in *P. angustifolia*, in *P. latifolia* males are distinguished from hermaphrodite individuals by the presence of an aborted stigma in the flowers they produce, in the left panel a hermaphrodite flower with a red arrow pointing to its functional stigma. In the right panel, a male flower with a red arrow pointing to the underdeveloped stigma.

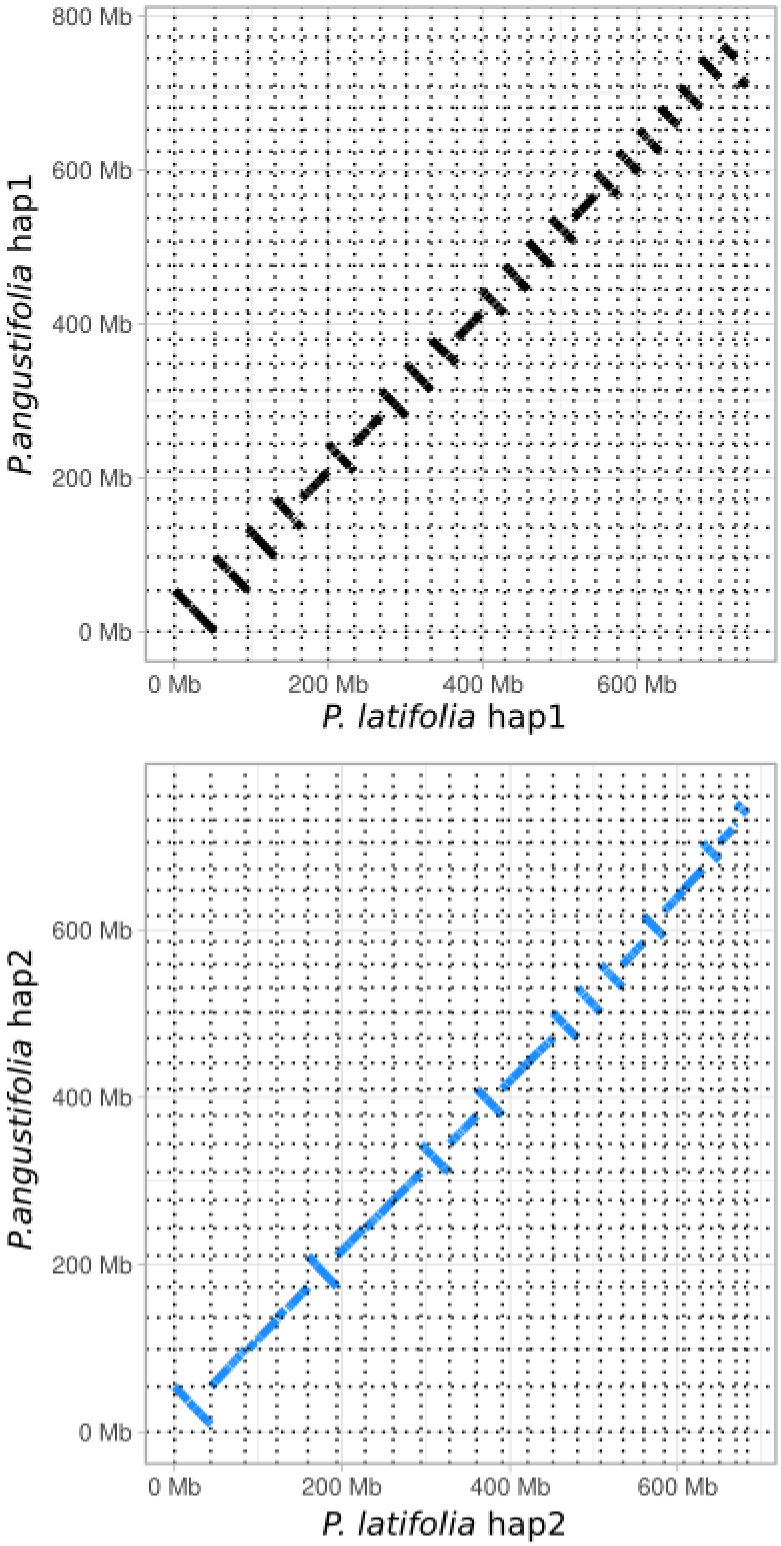


**Figure S9. Whole-genome alignment (WGA) between the male genome of *P. angustifolia* (040A11) and the newly sequenced male genome of *P. latifolia*.** The WGA was computed using minimap2 with the "-asm5" parameter. It reveals a high degree of synteny and contiguity between the two haplotype assemblies.

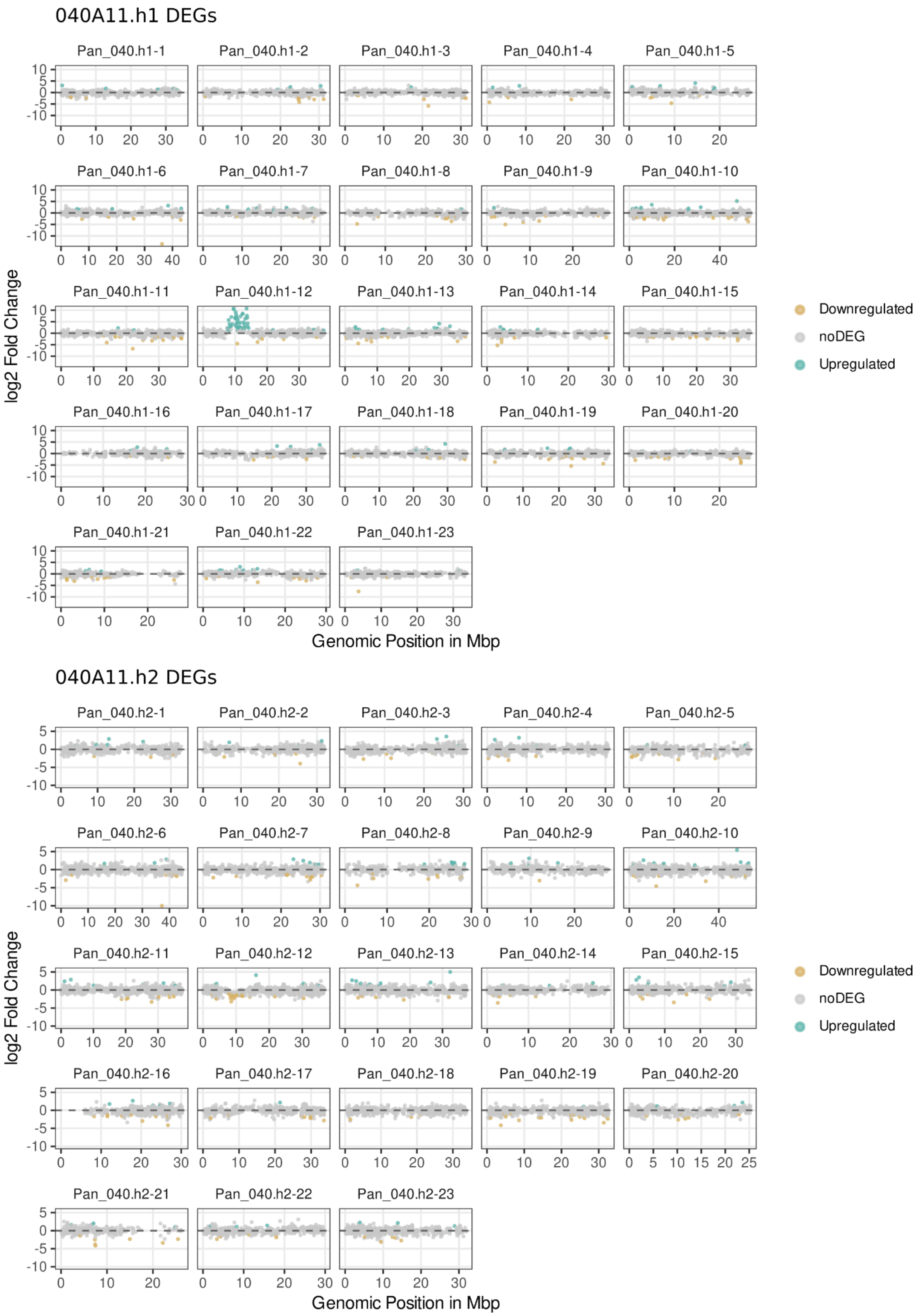


**Figure S11. Position along the genome of differentially expressed genes between males and hermaphrodites in flower buds in 040A11.** The upper panel shows gene expression data at haplotype 1, Y containing haplotype, and lower panel shows gene expression data at haplotype 2, X containing haplotype. Each point represents a gene with detectable expression, grey dots are non-differentially expressed genes whereas color dots represent differentially expressed genes, with turquoise color if they are more expressed in males and golden if they are more expressed in hermaphrodites.

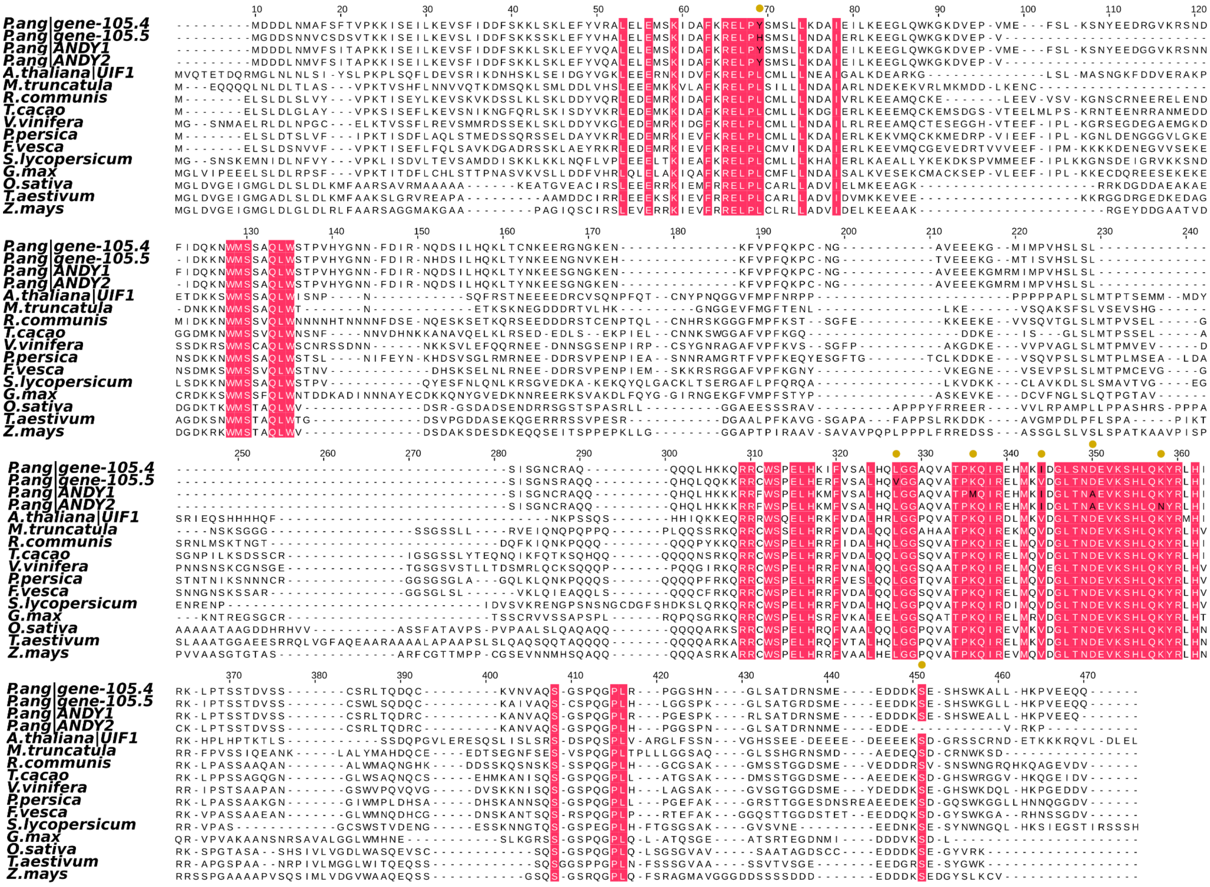


**Figure S12.** Reconstruction of the alignment reported by Moreau and colleagues (Moreau et al. 2016) of the amino acid sequences of *A. thaliana* UIF1 (AT4G37180.1), the 12 selected orthologous proteins: *Medicago truncatula* XP_003611799.1, *Ricinus communis* XP_002522328.1, *Theobroma cacao* XP_007047162.1, *Vitis vinifera* XP_002264629.2, *Prunus persica* XP_007204391.1, *Fragaria vesca* XP_004288067.1, *Solanum lycopersicum* XP_004233639.1, *Glycine max* XP_003528384.1, *Oryza sativa* Os03g0764600, *Brachypodium distachyon* XP_003558989.1, *Triticum aestivum* AEV91172.1 and *Zea mays* NP_001146647.1. The first two lines correspond to the predicted tandem homologous sequences in chromosome 14 of *P. angustifolia* and the next two to the additional male-specific tandem ANDY1/2 proteins. Red squares represent the conserved amino acids as reported by Moreau and colleagues (Moreau et al. 2016) with yellow dots if any amino acid diverges in the *P. angustifolia* orthologs with the corresponding amino acid in black.

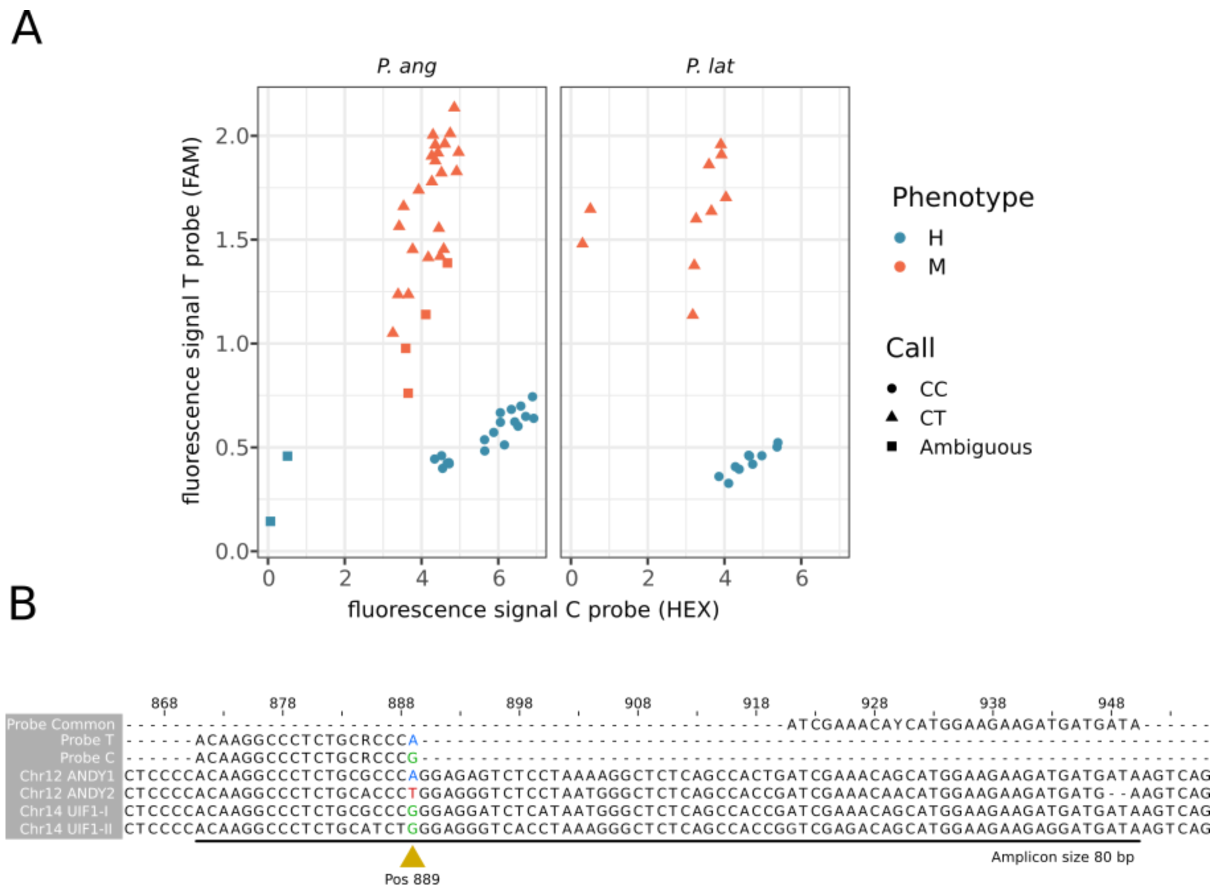


**Figure S13. Sex phenotypes are fully associated with SNP genotypes defined with the KASPar probe 889 in different *Phillyrea angustifolia* (Pang) and *P. latifolia* (Plat) accessions (see Table1). A)** The Left panel presents KASPar profiles observed for 10 *P. angustifolia* hermaphrodites and 16 males from a population located in Cadiz (Spain) and 9 hermaphrodites and 11 males from a population located in Sainte Lucie (France). The right panel presents KASPar profiles observed for 9 *P. latifolia* hermaphrodites and 10 males from a population located in Roussières (France). All hermaphrodites exhibit a (CC) signal (revealing the chr12 UIF1-I/II paralogs, acting as positive controls) and all males exhibit a (CT) signal (revealing the additional presence of ANDY2). **B)** Detailed alignment of the KASPar probe 889 at the four different *P. angustifolia* UIF1-like paralogs. SNP at position 889 distinguishes the presence of the Y-SDR in the sample. The total size of the amplicon was 80 bp.

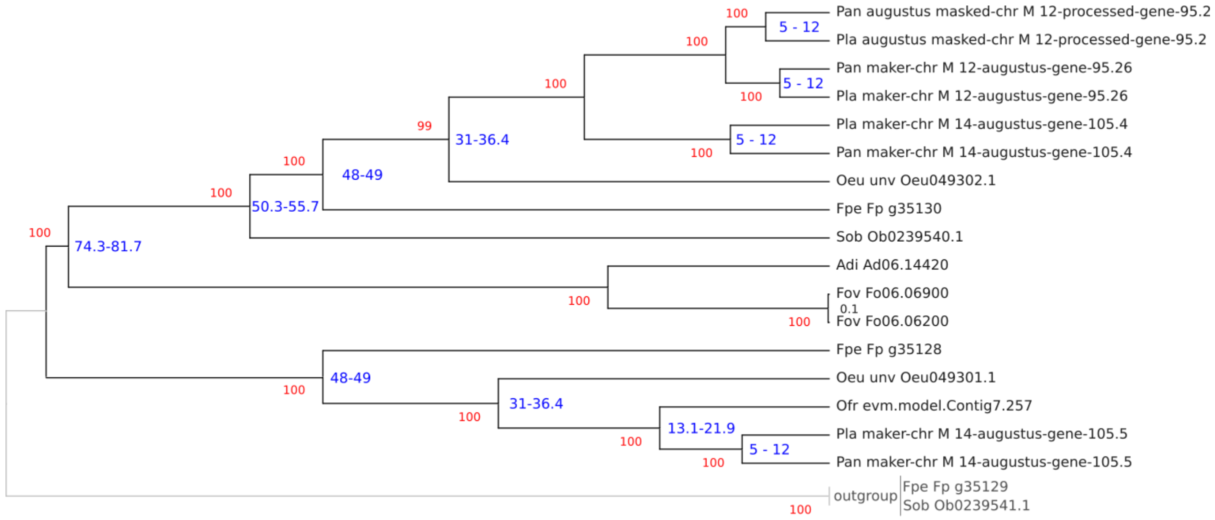


**Figure S14. Calibration tree used for MEGA's RelTime analysis.** This tree is scaffolded using the ML tree. Bootstrap support is displayed in red. In blue is reported the(Dupin et al. 2022) of each speciation event at each node as reported previously (Dupin et al. 2022; Dupin et al. 2024).

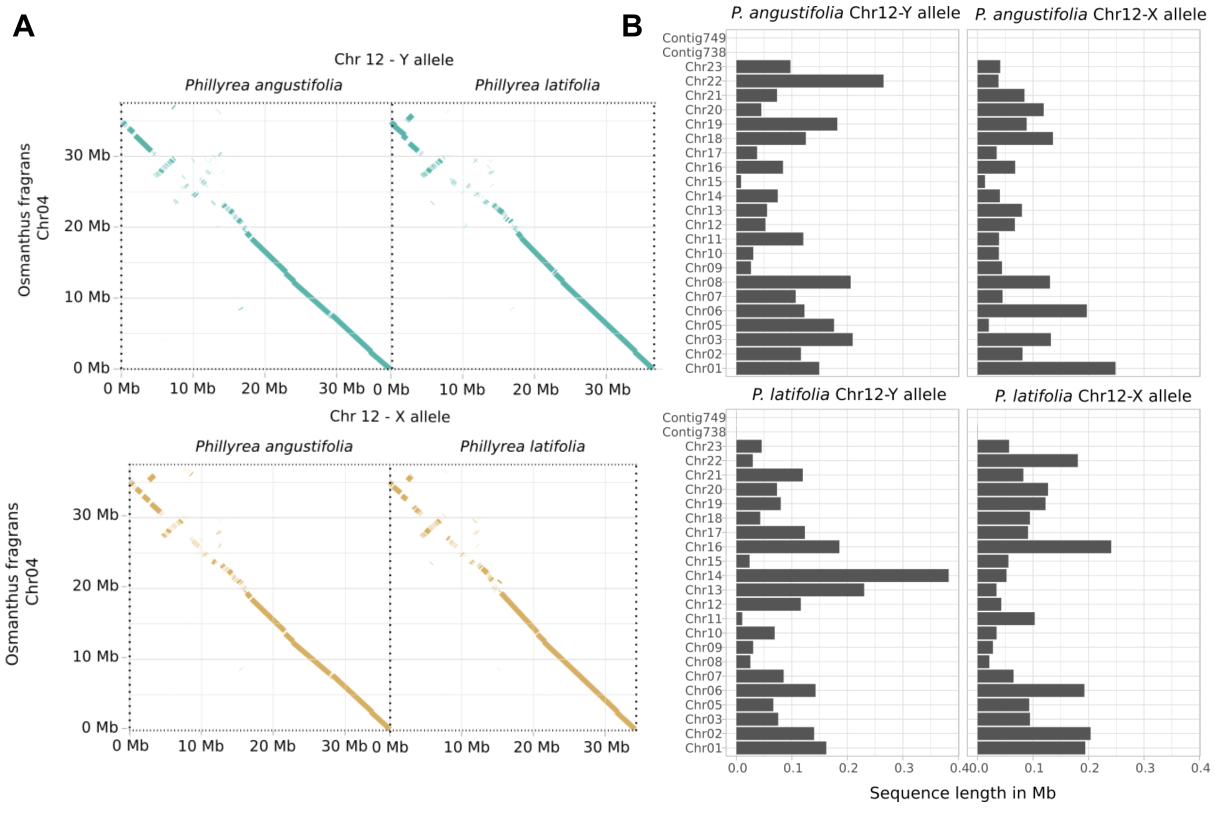


**Figure S15. Whole-genome alignment (WGA) between the haplotypes of the male *P. agustifolia* (040A11), the male of *P. latifolia* vs the primary genome of the publicly available male genome of *O. fragrans*.** A) WGA of Chr04 and X and Y alleles of both Phillyrea species. B) Total cumulative length of rest of alignments of *O. fragrans* against X and Y alleles of both species of Phillyrea excluding Chr 04.

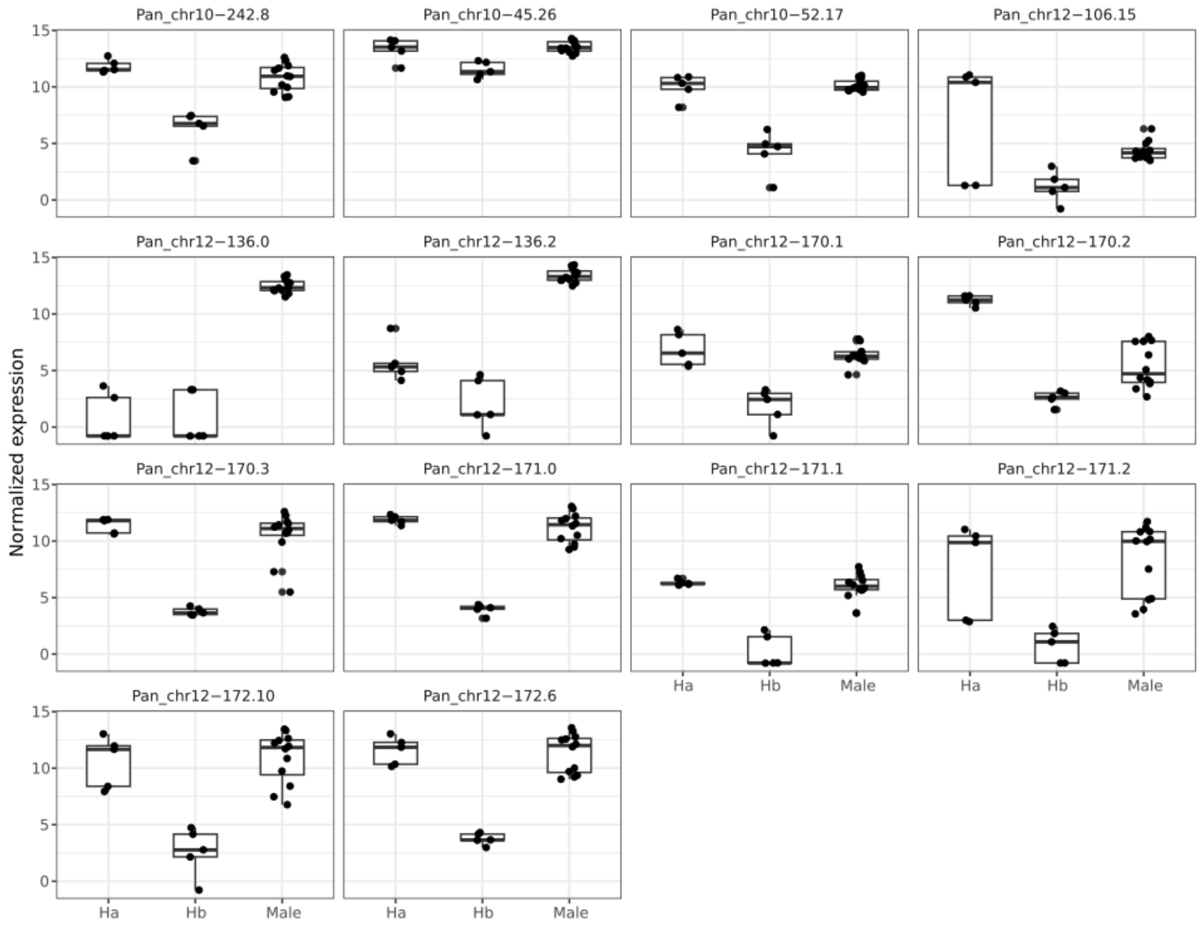


**Figure S16. Normalized expression counts of *Ole e1* paralogs in *P. angustifolia* in [Ha], [Hb] and [Male] individuals during flower development.** The expression of these 16 paralogs of Ole e1 during flower development more closely resemble those observed in hermaphroditic Ha individualism with the exception of Pan_chr12-106.15 and Pan_chr12-170.2, which exhibit uniquely high expression levels in Ha individuals. Males show high expression of the two Y-SDR paralogs, Pan_chr12-136.0 and Pan_chr12.136.2 which are not present in hermaphrodite individuals. Despite this, we find some residual mapping in both hermaphrodites, probably due to the high sequence similarity between all paralogs.

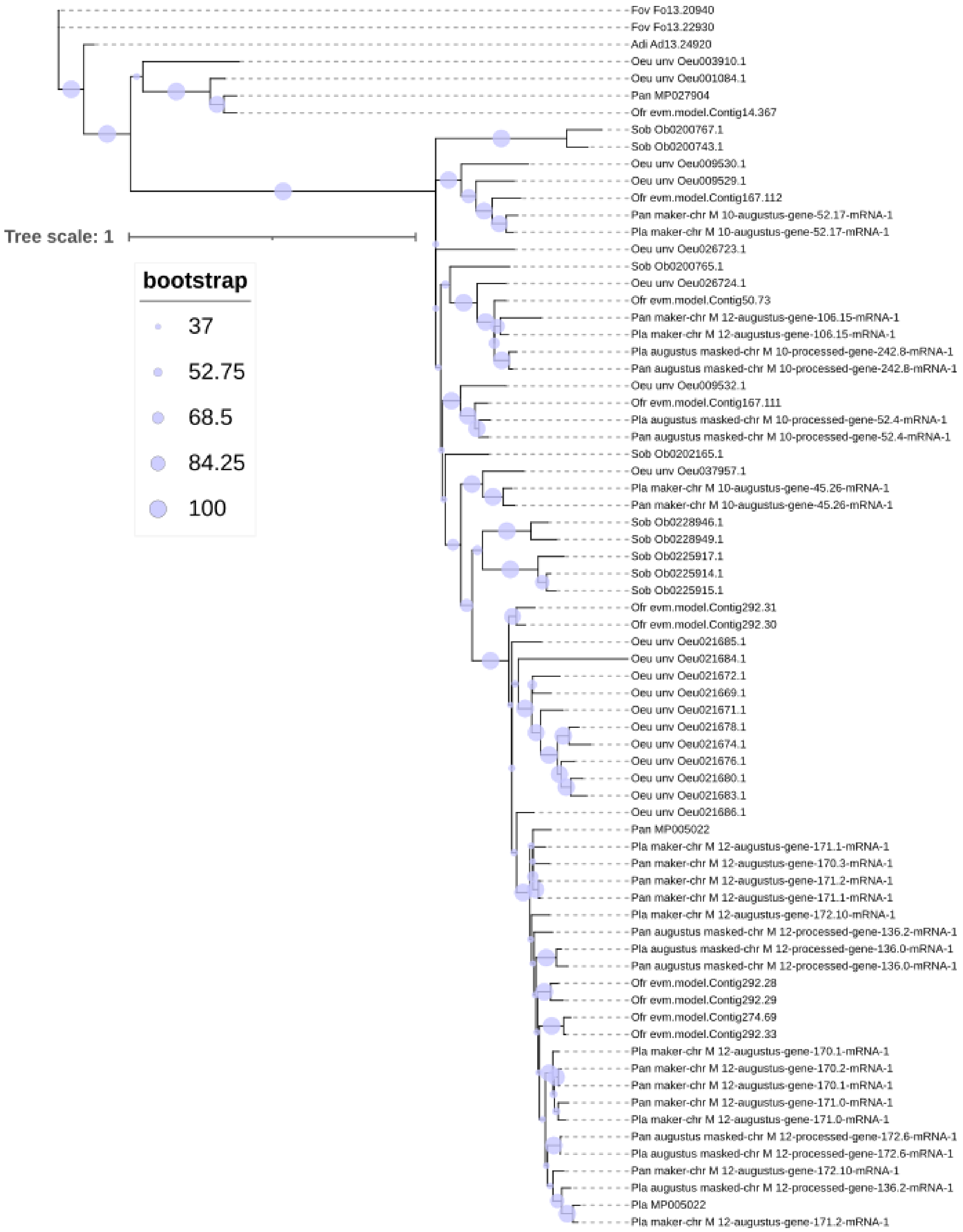


**Figure S17. Maximum-likelihood phylogenetic tree of the Ole e1 gene family**. This is the same tree presented in Fig. 6C, with all bootstrap support values and branch lengths displayed. Tip labels indicate the gene identifiers of the individual sequences included in the phylogenetic analysis.
